# Adaptative and ancient co-evolution of integrons with *Xanthomonas* genomes

**DOI:** 10.1101/2025.06.18.660453

**Authors:** Elena Colombi, Timothy M. Ghaly, Vaheesan Rajabal, Liam D.H. Elbourne, Michael Gillings, Sasha Tetu

**Affiliations:** School of Natural Sciences, Macquarie University, Sydney, NSW, 2109, Australia; ARC Centre of Excellence in Synthetic Biology, Sydney, NSW, 2109, Australia

**Keywords:** genome evolution, horizontal gene transfer, niche adaptation, plant-pathogen, mobile genetic elements

## Abstract

Integrons are genetic elements that facilitate gene acquisition. They have been extensively studied in clinical bacteria, but their evolutionary role in phytopathogens remains underexplored. Here, we analysed complete genomes of *Xanthomonas* species to investigate the origin, distribution, and functional dynamics of integrons in this genus. We found that 93% of genomes harboured integrons. The integron-integrase gene *intI* was predominantly located downstream of *ilvD*, indicating an ancestral acquisition of integrons, predating diversification within the genus. Phylogenetic analyses support vertical inheritance of *intI*, with the exception of rare horizontal gene transfer events, notably in *X. arboricola*. Despite their widespread presence, full-length *intI* genes and active integron platforms are only retained in some species, especially *X. campestris*, which shows high integron gene cassette variability and functional integron activity. In contrast, species such as *X. cissicola* and *X. phaseoli* exhibit widespread *intI* inactivation, likely occurring early in their divergence, leading to more stable cassette arrays and conserved integron-associated phenotypes. The number and diversity of genes within cassette arrays varied significantly by species and, to a lesser extent, by the ecological context of plant host cultivation. While most cassettes encoded proteins without a known function, those with annotated roles were associated with stress response mechanism, competitive exclusion, and plant-associated functions. Together, our findings demonstrate that integrons in *Xanthomonas* likely originated from a single ancient acquisition event, preceding genus-wide speciation, and have co-evolved with *Xanthomonas* pathovars as they adapted to distinct plant hosts.

**IMPACT STATEMENT:** This study provides the first comprehensive genus-wide analysis of integron evolution and dynamics in *Xanthomonas*, a globally distributed and versatile plant pathogen. Integrons are genetic platforms that allow horizontal transfer of genes. They are well characterised in human pathogens where they mediate transfer of antibiotic resistance genes, but little is known about their role in plant associated bacteria. We showed that in *Xanthomonas* the integron platform was ancestrally acquired, yet integrons have undergone repeated lineage-specific inactivation events. Despite widespread erosion of integron activity, some species such as *X. campestris* maintain robust and functionally diverse integrons that continue to shape genome plasticity. Notably, high cassette diversity, combined with the presence of rare and often uncharacterized genes within these arrays (some potentially involved in environmental sensing or host interaction) suggest that integrons may serve as reservoirs of adaptive potential. Our findings reshape current views of integron function beyond antibiotic resistance and highlight their long-term role in microbial evolution, niche adaptation, and genome innovation in plant-associated bacteria.

**DATA SUMMARY:** The authors confirm all supporting data, code and protocols have been provided within the article or through supplementary data files. Accession numbers of the genomes analysed in this manuscript are listed in Table S1.

## INTRODUCTION

Integrons are genetic elements that facilitate horizontal gene transfer in Bacteria (Stokes & Hall, 1989), and Archaea (Ghaly *et al*., 2022). They are ancient structures that have been involved in the evolution of bacterial genomes for hundreds of millions of years (Rowe-Magnus & Mazel, 2001). Functional integrons are composed of an integron-integrase gene (*intI*), an integron recombination site (*attI*), a promoter that drives the expression of the integrase (P_int_) and generally also have a second promoter (P_c_), oriented for expression of gene cassette array proteins (Hall & Collis, 1995, Mazel *et al*., 1998, Escudero *et al*., 2018).

Gene cassettes are mobile, non-replicating elements, which generally consist of an open reading frame (ORF) and an *attC* site. They exist as circular molecules when excised from a cassette array. Although cassettes generally consist of a single ORF, cassettes that have two ORFs or ORF-less cassettes with promoter activity have been observed (Michael & Labbate, 2010, Tansirichaiya *et al*., 2019, Blanco *et al*., 2024). The tyrosine recombinase IntI can capture circular cassettes and integrate them into the genome by mediating the recombination between *attC* and *attI* sites, forming arrays of 1 to > 200 gene cassettes. As well as mediating the insertion of cassettes, IntI can also excise cassettes from an array. Cassette arrays lacking an integrase are called CALIN (<u>c</u>lusters of *attC* sites lacking <u>in</u>tegron-integrases) (Cury *et al*., 2016), and can be mobilized *in trans* by a functional IntI.

Technically, integrons themselves are not mobile, whereas their gene cassette cargo is. Integron activity can facilitate horizontal gene cassette transfer between prokaryotes, contributing to the rapid evolution and genetic diversity of Bacteria and Archaea (Escudero *et al*., 2018). Because DNA damage and nutrient starvation induce activity of the integron integrase via bacterial SOS and stringent responses (Guerin *et al*., 2009, Cambray *et al*., 2011, Strugeon *et al*., 2016), acquisition of novel genetic functions is preferentially triggered during periods of environmental challenge. This provides a mechanism for adaptive innovation when survival pressures are highest.

Integrons have been extensively studied because of their role in the accumulation and dissemination of antibiotic resistance genes in clinical isolates (Gillings *et al*., 2008, Hipólito *et al*., 2023). However, integrons are also common in other environments such as marine environments (Pereira *et al*., 2020), soil (Ghaly *et al*., 2019), and the plant rhizoplane (Qi *et al*., 2024, Rajabal *et al*., 2024). In these environments, the majority of gene cassettes remain functionally uncharacterised, but predicted functions encode extensive functional diversity including traits involved in microbe-host interactions (Gillings, 2014, Escudero *et al*., 2018, Ghaly *et al*., 2020).

*Xanthomonas* is a genus of Gram-negative bacteria in the class Gammaproteobacteria that contains pathogenic strains able to cause disease in more than 400 different plants. In common with other genera of plant pathogens, *Xanthomonas* also contains strains associated with plants, that do not cause disease symptoms (Vauterin *et al*., 1996, Essakhi *et al*., 2015, Garita-Cambronero *et al*., 2017). *Xanthomonas* is a large genus that encompasses more than 35 species, which are subdivided into pathovars (Timilsina *et al*., 2020).

Integrons have been described in some *Xanthomonas* isolates (Gillings *et al*., 2005, Barionovi & Scortichini, 2006, Barionovi & Scortichini, 2008). Using PCR with site-specific primers annealing to the proximal part of *intI* and to the *attI* region, Gillings and colleagues screened 32 *Xanthomonas* strains representing 12 pathovars, and found that all the strains analysed had an integron. In all cases, *intI* was integrated downstream of the acid dehydratase gene, *ilvD*, however, the majority of integrase genes were predicted to be inactivated by frameshifts, stop codons, or large deletions. Groups of strains with the same deletions or stop codons/frameshifts in *intI* usually contained identical arrays of genes. There was no evidence of integrons generating diversity within pathovars but strong evidence for diversity between pathovars. With some minor exceptions, individual pathovars had distinct proximal gene cassette arrays, and every cassette identified was found only in one pathovar.

Complete genome sequences for hundreds of *Xanthomonas* isolates are now available, allowing for a systematic and detailed investigation of integrons in these important plant-associated bacteria. Here, we have examined the presence of integrons in all publicly available complete genomes of *Xanthomonas*, significantly expanding our knowledge of these elements, and showing how integrons have been a dynamic component during *Xanthomonas* genome evolution.

## METHODS

### *Xanthomonas* classification and phylogeny

Complete genomes of *Xanthomonas* were downloaded from the National Center for Biotechnology Information (NCBI) Database (NCBI txid: 338) (708 genomes, last accessed September 2024) (Table S1). Taxonomic classifications of the genomes was based on the Genome Taxonomy Database (GTDB) (Parks *et al*., 2022) release 2.2.0 using GTDB-Tk v2.4.0 (Chaumeil *et al*., 2022) (Table S1). The command classify_wf was used with default settings. GTDB-Tk first aligns 120 single-copy phylogenetic marker genes, then classifies each genome based on its placement into domain-specific reference trees (built from 47,899 prokaryote genomes), its relative evolutionary divergence, and average nucleotide identity to reference genomes in the GTDB. Two genomes (GCA_041228075.1 and GCA_002240395.1) did not belong to the *Xanthomonas* genus (*Luteibacter* and *Luteimonas*_D, respectively) and were removed from the dataset. The GTDB-Tk classification system does not strictly adhere to the traditional nomenclature of *Xanthomonas* strains, sometimes reassigning species. We use GTDB-Tk classification in order to use a standardised microbial taxonomy method based on genome phylogeny, and not to attempt to re-classify or rename isolates.

Phylogeny of the *Xanthomonas* genus was inferred using on the alignment of the 120 marker genes produced with the GTDB-Tk command classify_wf using GTDB-Tk command infer (default setting with --gamma) which uses FastTree 2.1 (Price *et al*., 2010) to build an approximately-maximum-likelihood phylogenetic tree. The tree was rooted with GCA_041228075.1 (*Luteibacter*) then the tip was removed with the R package ape 5.8-1 (Paradis & Schliep, 2019). Single-species phylogeny was built using Realphy (Bertels *et al*., 2014) which maps genomes to a series of reference genomes via bowtie2 2.0.0-beta7 (Langmead & Salzberg, 2012). From these, multiple sequence alignments are constructed and used to infer phylogenetic trees via PhyML 3.1 (Langmead & Salzberg, 2012). Trees are visualised using the ggtree 3.16 (Yu *et al*., 2017) and ggtreeExtra 2.3 (Xu *et al*., 2021) R packages.

### Analysis of integrons

Integrons were identified with IntegronFinder 2.0.5 (Néron *et al*., 2022) (parameters: --local-max --gbk --promoter-attI) (Table S1). IntegronFinder 2.0.5 was run with default --calin-threshold (default: 2), putative CALINs were identified if they carried at least 2 *attC* sites. To determine the genomic location of the integrons, identified integron sequences were extracted together with 5 kb of flanking regions upstream and downstream (fasta files deposited at https://github.com/EC-MQ/Xanthomonas_integrons) and annotated with Bakta 1.11.3 (Schwengers *et al*., 2021). Taxonomic origins of integron gene cassette recombination sites (*attC*s) were predicted using the script attC-taxa.sh (Ghaly *et al*., 2021, Ghaly *et al*., 2022). The *attC*-taxa pipeline (https://github.com/timghaly/attC-taxa) can detect *attC* sites that are conserved among one of 11 bacterial taxa, comprising: Alteromonadales, Methylococcales, Oceanospirillales, Pseudomonadales, Vibrionales, Xanthomonadales, Acidobacteria, Cyanobacteria, Deltaproteobacteria, Planctomycetes and Spirochaetes.

*IntI* sequences identified by Integron Finder were aligned using PRANK (Löytynoja, 2013) run with default settings, the alignment was stripped of gaps with the goalign ‘clean sites’ command (Lemoine & Gascuel, 2021) and the tree inferred with FastTree 2.1 (Price *et al*., 2010) using the generalized time-reversible model (--gtr).

We used multiple approaches to annotate the cassette-encoded proteins. Proteins were functionally annotated with eggNOG-mapper v2 (Huerta-Cepas *et al*., 2019, Cantalapiedra *et al*., 2021), executed in DIAMOND (Buchfink *et al*., 2015) mode. Additionally, Foldseek (Van Kempen *et al*., 2024) was used to create a protein structural database with the ProstT5 protein language model (Heinzinger *et al*., 2024). The database was then used to perform structural alignment against the SwissProt database using a Foldseek search. Foldseek convertalis was used to convert the alignment database in a tab-delimited output, only functional annotations against the Swiss-Prot database with a coverage of 80% and an e-value < 0.001 were considered.

To identify genes involved in functions of ecological relevance (trace gas oxidation, carbon cycling, nitrogen cycling, sulphur cycling, phosphorus cycling, iron cycling, plant-microbe interactions, and osmotic stress tolerance) we used the *EcoFoldDB-annotate* pipeline (Ghaly *et al*., 2025).

Plant growth-promoting traits were identified using the Plant Growth-Promoting Traits Prediction tool (PGPT-Pred) (Ashrafi *et al*., 2022). SignalP V6.0 (Almagro Armenteros *et al*., 2019) was used for signal peptide predictions as a marker for identifying transmembrane or secreted gene products. DefenseFinder (Tesson *et al*., 2022) was used to detect known anti-phage systems.

Unique gene orthologues encoded by the cassettes and their distributions were identified with Proteinortho 6.0.22 (Klemm *et al*., 2023) run with the -singles option.

To detect genes encoding type III secretion effectors (T3SE), we used a database of 66 previously characterized 66 *Xanthomonas* T3SE retrieved from the EuroXanth platform (Costa *et al*., 2024) (Biessy *et al*., 2025). T3SE in the cassettes were identified with Proteinortho v6.0.22 (Klemm *et al*., 2023) by querying the curated effectors database against the amino acid sequences of the cassette-encoded genes.

Phylogenetic signal in the number of *attC* sites across genomes was first assessed with Pagel’s λ tests using the R package caper 1.0.3 (Orme *et al*., 2013). To evaluate the influence of cultivation type on *attC* site number while accounting for shared evolutionary history, phylogenetic generalized least squares regression was conducted using the R package nlme 3.1 (Pinheiro *et al*., 2017).

### Horizontal gene transfer events between species

Alfy 1.0.5 (Domazet-Lošo & Haubold, 2011) was used to guide the detection of highly similar cassettes present in the integron arrays. Alfy was run with the -M option, and selecting only matches with a *P* value <0.05 within a sliding window of 100 bp. Only arrays sharing closest homology (recombination), with at least 10% of their array sequence, were considered. The output of this was further filtered, using average nucleotide identity comparisons to assess likelihood that shared cassettes represented probable horizontal gene transfer (HGT) events. FastANI 1.33 (Jain *et al*., 2018) was run between the arrays and the whole genomes of the strains. We classified an event as HGT when the ANI identity between arrays exceeded 96%, the ANI between the genomes harbouring them was below 95%, and the array ANI was at least 2% higher than the genome ANI. Cytoscape (Shannon *et al*., 2003) was then used to visualise the numbers of cassettes transferred between *Xanthomonas* species.

## RESULTS AND DISCUSSION

### The *Xanthomonas* integron platform is ancestral

*Xanthomonas* complete genomes classified based on the Genome Taxonomy Database (GTDB) as “*Xanthomonas*” (n=629) and “*Xanthomonas_A*” (n=78), known also as *Xanthomonas* group 2 and 1, respectively (Pena *et al*., 2024) were analysed for integron features (Table S1). We retained the genomes of *Xanthomonas_A,* even though they are classified as a different genus by GTDB-Tk, because from a plant pathology perspective they are still considered *Xanthomonas*, and encompass *X. translucens,* an important causal agent of diseases in cereal crops and forage grasses (Sapkota *et al*., 2020).

Using IntegronFinder 2.0.5 (Néron *et al*., 2022), we identified integrons, In0 (integron-integrases that lack cassettes), or CALINs (clusters of *attC* sites lacking integron-integrases) in 93% of the genomes (657 genomes) (Figure 1, Table S1). *X. albilineans* (n=6, group 1) and *X. fragariae* (n=6, group 2) were the only species with more than 2 genome sequences available that did not harbour either an *intI* or *attC* sites. However, integrons have been previously detected in some isolates of *X. fragariae* via PCR amplification (Barionovi & Scortichini, 2006).

**Figure 1.**
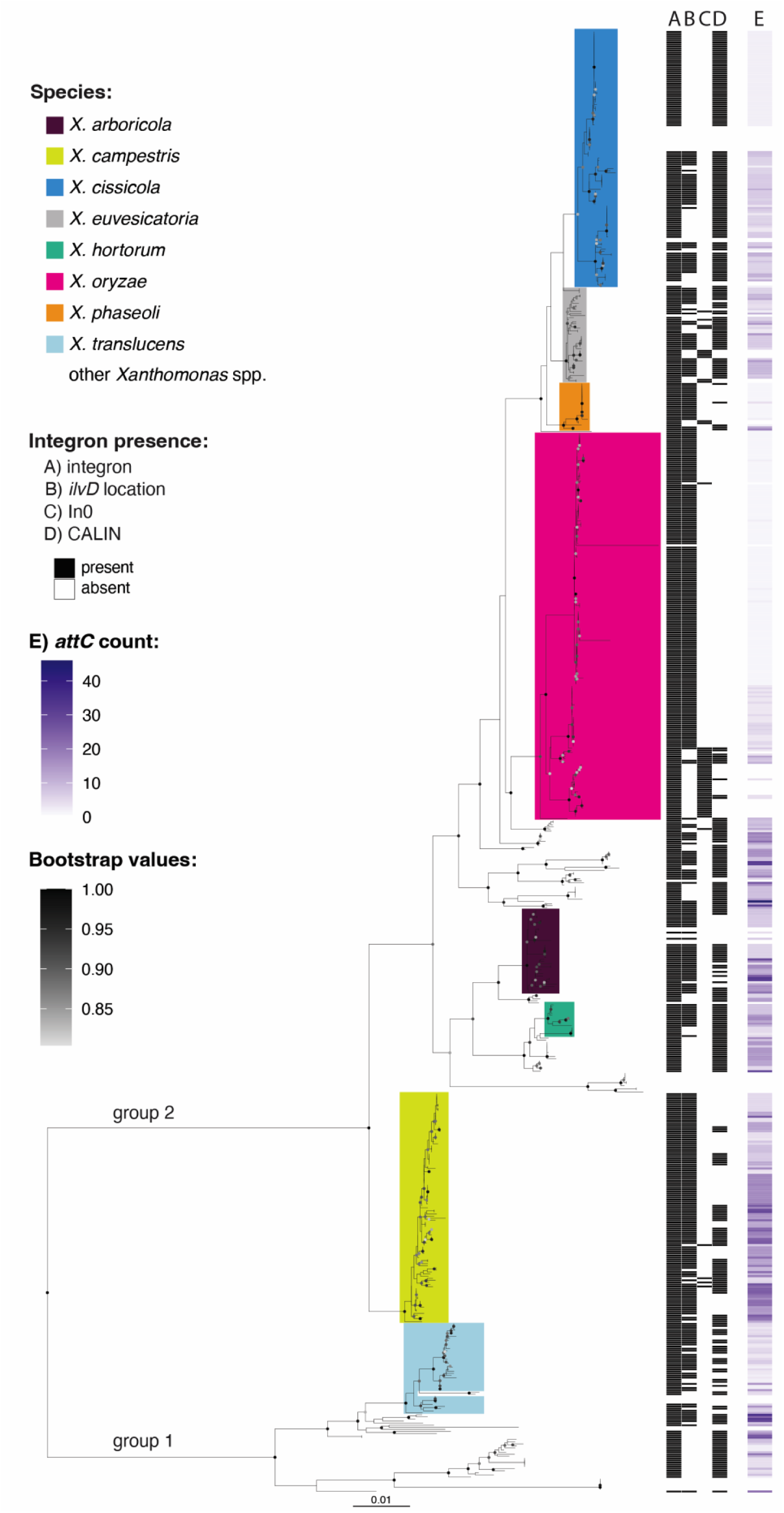
Distribution of integron platforms in *Xanthomonas* groups 1 and 2. The maximum likelihood phylogenetic tree is based on a concatenated alignment of 120 top-ranked marker proteins. The tree was rooted at the midpoint. Tips are labelled with species names assigned with Genome Taxonomy Database phylogeny. Coloured clades show species with more than 10 genomes available for analysis. A) Integrons were classified as present if the isolates carried a functional integron, *intI* only or an *attC* array only, B) Indicates the presence of a complete integron or *attC* sites in the *ilvD* locus, C) indicates the presence of an *intI* lacking cassettes (In0), D) indicates the presence of a CALINs (cluster of *attC* sites) not in the *ilvD* locus. E) indicates the number of *attC* sites (number of cassettes) in the genome. Only bootstrap values 0.80 are shown. Scale bar indicates substitutions per site.

Because integrons are widespread in *Xanthomonas*, our first question was whether the acquisition of the integron module was ancestral. The integron-integrase encoding gene *intI* was integrated downstream *ilvD* in all genomes, with the two exceptions being isolates of *X. oryzae* and *X. cucurbitae*. Of the 187 *oryzae* isolates examined, *X. oryzae* pv. *oryzicola* (26 genomes) harbour a truncated *intI* (561 bp) that was not integrated downstream *ilvD.* However, the integron locus (*intI* and cassette array) is surrounded by transposases, which could indicate that the integron was moved via transposition, or could be a result of assembly errors which are common in regions flanked by insertion sequences. In all seven *X. cucurbitae* genomes, *intI* was not integrated downstream *ilvD.* However, downstream of *ilvD* there are two genes without recognizable *attCs* which are encoded in the other integrons (including a DUF1488 encoding gene which occurs in multiple species).

The different genomic locations of *intI* could be explained by either within-genome transposition or recombination (*intI* was initially located downstream *ilvD* and then moved), or by an independent acquisition of *intI* from another source which was integrated in a different locus. To test this, we constructed the phylogeny of all the *Xanthomonas intIs* (Figure S1, Table S2). If *intIs* located elsewhere from *ilvD* were acquired independently, they would likely have a different evolutionary history, and form separate clades in the *intI* phylogeny.

The phylogeny of *intI* clusters this gene within species, generally being independent of the integration locus. *X. oryzae intIs* cluster together, and the *X. cucurbitae intIs* form a clade with pv. *phaseoli* (Figure S1). The only notable exception is *X. arboricola*. *X. arboricola* strains can carry one of two variant *intIs*. In two strains, *intI* clusters closest to the *X. campestris intIs* and are truncated and likely not functional. Phylogenetically close *X. arboricola* strains do not harbour an *intI* at all. The other *X. arboricola* strains carry *intI* (either full-length or with an early stop codon) that are more distantly related to all other *intIs*. These more distantly related *intI* could have been acquired from a source outside *Xanthomonas*. We compared the *ilvD*-*intI* locus sequence in two representative *X*. *arboricola* strains harbouring the two *intI* variants (GCA_018141705.1 and GCA_905367745.1). While the *ilvD* genes share 97.1% average nucleotide identity (ANI), the *intIs* share 68.2% ANI, this discrepancy in ANI values between two adjacent genes suggests a recombination event (Ansari & Didelot, 2014), with abrupt shifts in ANI at gene boundaries a common indicator of a recombination breakpoint.

We used two full length *intI* variants (one belonging to *X. arboricola*, one to *X. campestris*) as a query in a BLASTn search on complete genomes in NCBI excluding *Xanthomonas* (taxid:338). The two closest *intI*s belonged to *Lysobacter* sp. CECT 30171 (locus tag: LYB30171_00805, 93% coverage and 74% identity against *X. campestris intI*) and *[Pseudomonas] boreopolis* strain GO2 (locus tag: M3M27_18655, 92% coverage and 91% identity against *X. arboricola intI*) (both genera in the Xanthomonadaceae family). Curiously, in isolate GO2, *intI* was also located downstream of *ilvD*. We constructed a phylogenetic tree using *Xanthomonas intI* sequences longer than 900 bp and included the *intIs* of CECT 30171 and GO2, using *Vibrio* sp. SCSIO 43136 *intI* as an outgroup (Figure 2). The *intI* variants in *X. arboricola* were more closely related to the *intI* in GO2 than the other *Xanthomonas intIs*. *ilvD* phylogeny instead clusters together all *X. arboricola* isolates within the *Xanthomonas* group 2, and GO2 as basal both to *Xanthomonas* group 1 and 2 isolates (Figure S2). This suggests that, in those *X*. *arboricola* strains, *intI* was acquired via horizontal gene transfer (HGT) from another member of the Xanthomonadaceae family.

**Figure 2.**
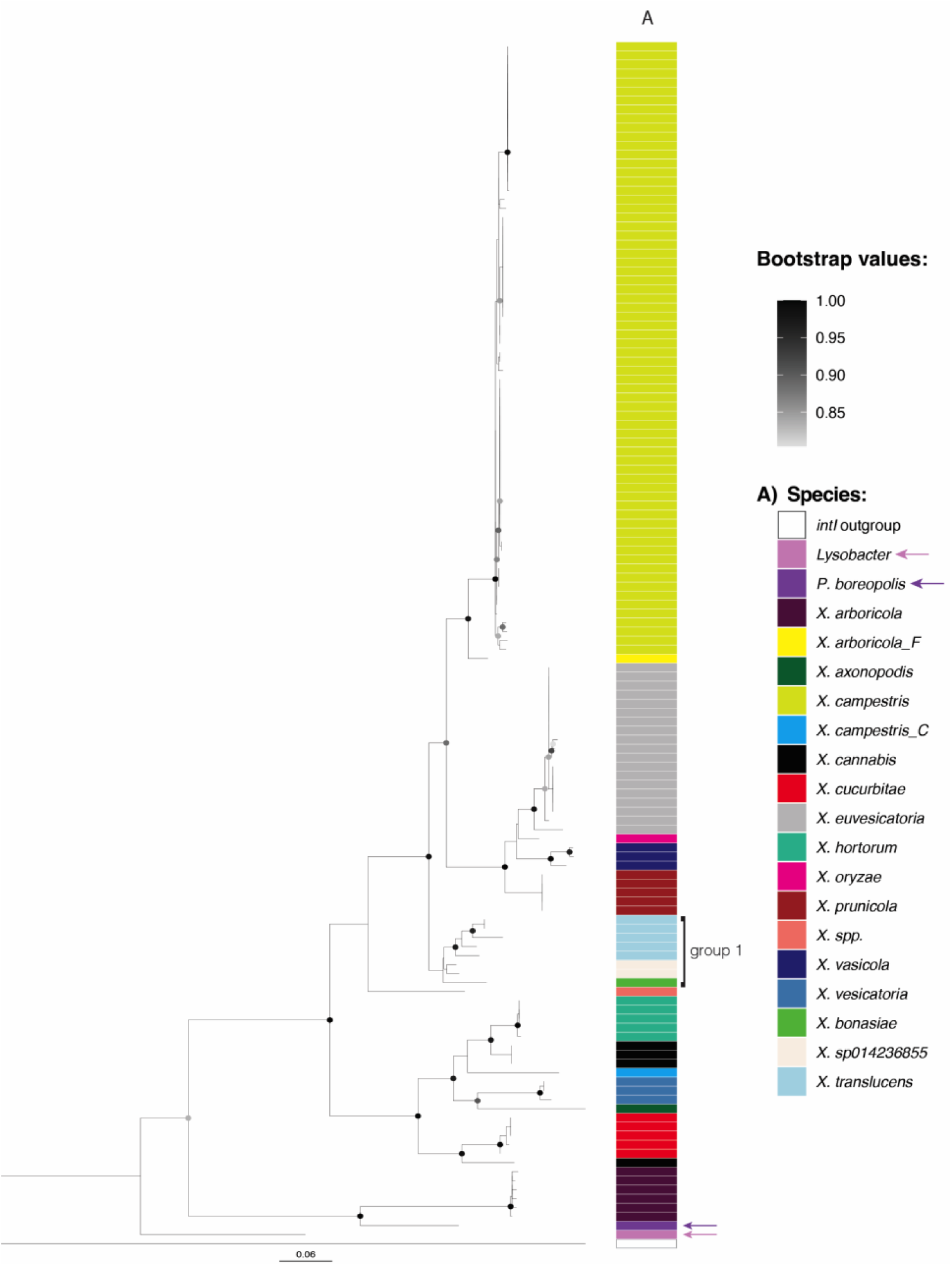
Phylogeny of *intI*. Sequences of *intI* longer than 900 bp were aligned with PRANK, stripped of gaps and used to infer the tree with FastTree. *intI* from *Vibrio* sp. SCSIO 43136 (locus tag: J4N39_08275) was used as an outgroup. *intI* from group 1 *Xanthomonas* are indicated on the side of the plot, arrows indicate *intI*s of *Lysobacter* sp. CECT 30171 and [*Pseudomonas*] *boreopolis* strain GO2. The scale bar indicates substitutions per site. Only bootstrap values 0.80 are shown.

The phylogeny of *ilvD* and *intI* is incongruent, with the *ilvDs* of *Xanthomonas* group 1 forming a separate group in comparison to group 2 (Figure S2), while the group 1 *intIs* are nested within the most common *Xanthomonas intIs* (Figure 2). Genetic exchange between a restricted number of strains of group 2 and the entire group 1 clade has been reported (Pena *et al*., 2024), therefore it is plausible that horizontal transfer of an *intI* gene from a group 2 *Xanthomonas* species occurred during the early evolutionary history of group 1 *Xanthomonas*.

Together, this suggests that the acquisition of the integron platform was ancestral to the *Xanthomonas* genus, and then *intI* diversified with the species. The exceptions are some *X. arboricola* strains that could have recruited *intI* from another source, while group 1 *Xanthomonas* could have acquired *intI* from group 2 *Xanthomonas* in one single gene flow event.

Additionally, approximately 50% of genomes harbor CALINs, here defined as clusters of at least two *attC* sites, in loci other than the canonical *ilvD* region and in the absence of a proximal *intI* gene. In genomes where *intI* is retained, this gene is consistently integrated downstream of *ilvD,* while these CALINs show spatial separation, being located in other regions of the genome. Among the species represented by more than 10 genomes, CALINs not adjacent to *ilvD* were identified at seven distinct chromosomal loci. In *X. arboricola*, *X*. *campestris*, *X*. *cissicola*, *X*. *euvesicatoria*, and *X*. *hortorum*, CALINs consistently occurred adjacent to *lamG* and/or *secF*. In *X. cissicola*, additional CALINs were located near *recD* and a gene encoding a NAD(P)/FAD-dependent oxidoreductase. In *X. oryzae*, two strains possessed CALINs adjacent to an outer membrane protein gene, while in *X. translucens*, CALINs were observed at two distinct genomic positions. One *X. campestris* strain (GCA_028749605.1) harboured a CALIN on a conjugative plasmid. The relatively conserved positioning of CALINs across species supports the hypothesis of ancestral integron activity. This contrasts with experimental data from *Escherichia coli*, where cassette libraries integrated via *attG* sites (consensus recombination sequences that are not *attC* or *attI*) exhibited a broad distribution across numerous genomic loci, indicating that *de novo* integration is typically less site-specific (Loot *et al*., 2024).

### Acquisition of the integron platform is ancestral but its activity is progressively lost

To investigate integron integrase activity and the complement of integron gene cassettes in *Xanthomonas* genomes, we focused on species which had more than 10 genomes in our dataset: *X*. *campestris* (n=111) (Figure 3), *X*. *cissicola* (n= 124) (Figure 4), *X*. *arboricola* (n=41) (Figure S3), *X*. *translucens* (n=46) (Figure S4), *X*. *euvesicatoria* (n=46) (Figure S5), *X*. *oryzae* (n=161) (Figure S6), *X*. *phaseoli* (n=23) (Figure S7), *X*. *hortorum* (n=12) (Figure S8). A detailed description of the integrons in these eight species is reported in the Supplementary Results Section.

**Figure 3.**
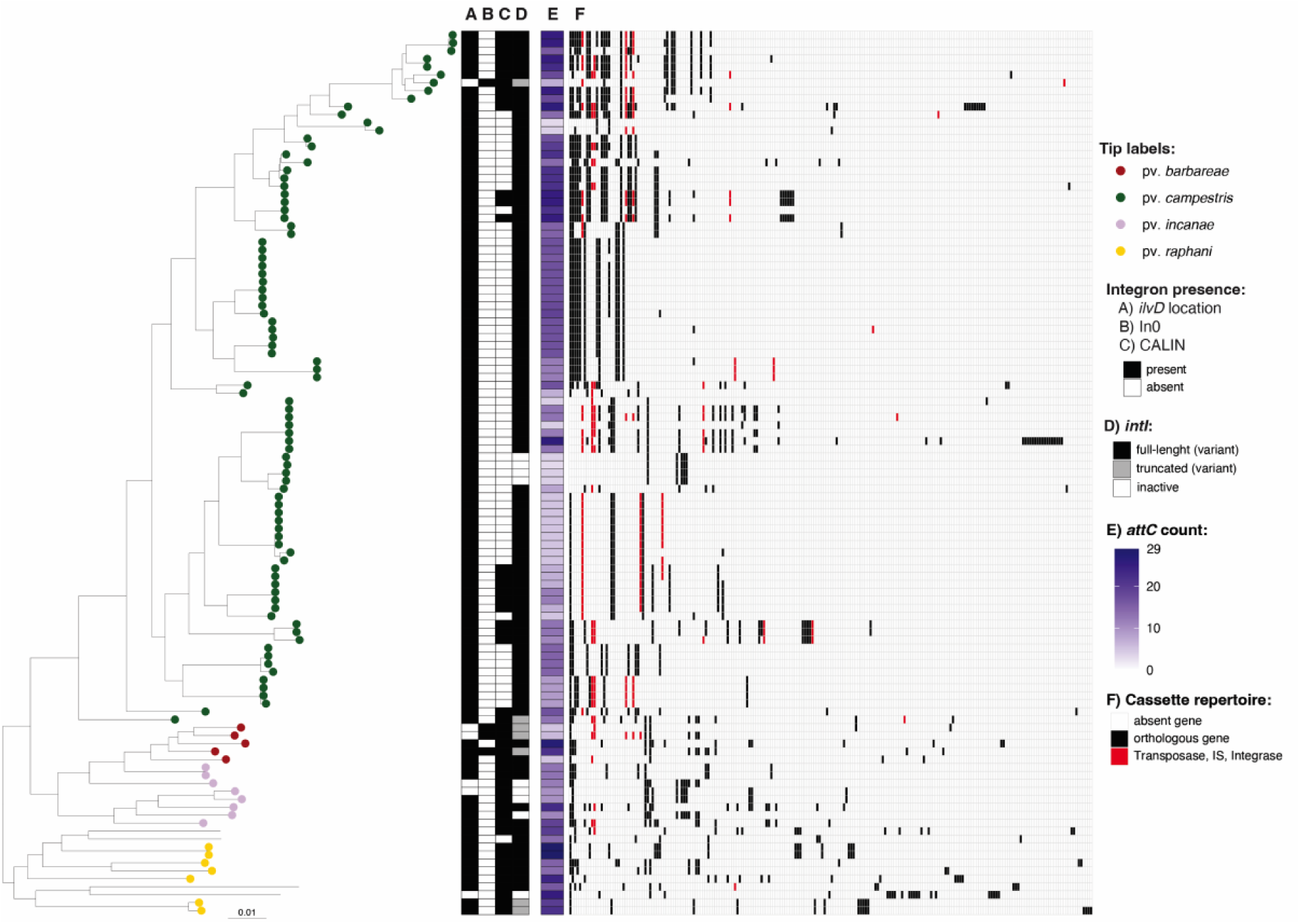
Integrons in *X. campestris*. The phylogeny of *X. campestris* was built using Realphy, using GCA_000007145.1 as a reference genome. The tree was rooted with GCA_000972745 (*X. arboricola*) then the tip was removed from the tree. Tip labels show pathovars, which were assigned from the literature search (Table S3). The scale bar indicates substitutions per site. A) Indicates the presence of a complete integron or *attC* sites in the *ilvD* locus, B) indicates the presence of an *intI* lacking cassettes (In0), C) indicates the presence of a CALINs (cluster of *attC* sites) not in the *ilvD* locus, D) indicates whether IntI is predicted to be functional (full length) (Table S3). E) indicates the number of *attC* sites (number of cassettes) in the genome. Panel F) represents the distribution of orthologous genes among all cassettes carried by the corresponding isolate inferred with Proteinortho. Orthologous genes are listed in decreasing order based on the number of strains within the species that carry them.

**Figure 4.**
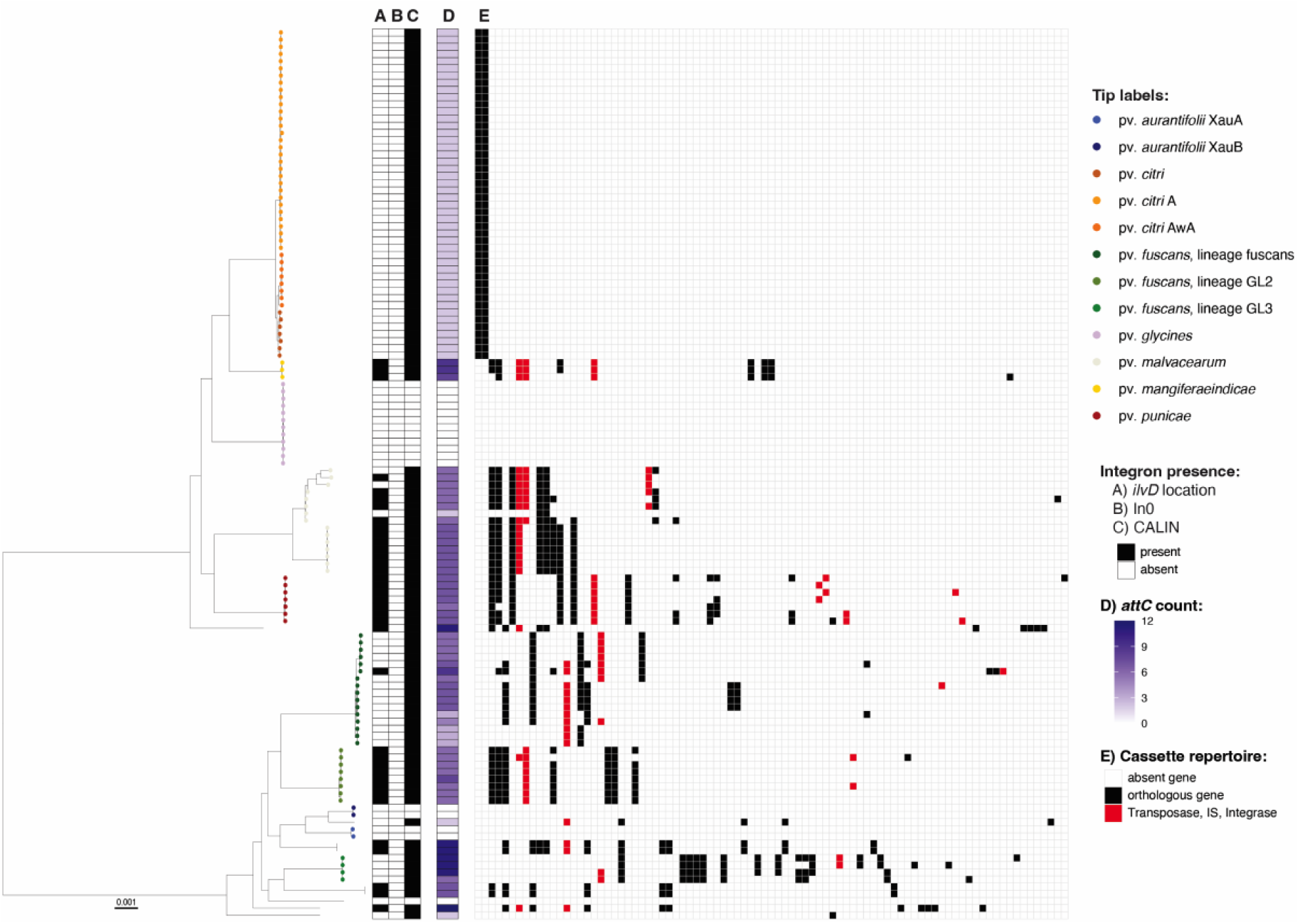
Integrons in *X. cissicola*. The phylogeny of *X. cissicola* was built using Realphy, using GCA_000007165.1 as a reference genome. The tree was rooted with GCA_000009165.1 (*X. euvesicatoria*) then the tip was removed from the tree. Tip labels show pathovars, which were assigned from the literature search (Table S3). The scale bar indicates substitutions per site. A) Indicates the presence of a complete integron or *attC* sites in the *ilvD* locus, B) indicates the presence of an *intI* lacking cassettes (In0), C) indicates the presence of a CALINs (cluster of *attC* sites) not in the *ilvD* locus, D) indicates the number of *attC* sites (number of cassettes) in the genome (Table S3). Panel E) represents the distribution of orthologous genes among the all the cassettes carried by the corresponding isolate inferred with Proteinortho. Orthologous genes are given in decreasing order based on the number of strains within the species that carry them.

Although integron associated sequences are common in *Xanthomonas* groups 1 and 2, most strains appear to have lost integron integrase activity (*intI* is either truncated or absent), and the distribution of functional integrons appears to be species specific. However, among the species analysed, *X. campestris* stands out for maintaining a largely functional integron platform (Figure 3). A high proportion (85.6%) of its genomes carry a full-length *intI* gene, and integrons are consistently flanked by extensive cassette arrays (ranging from 3 to 29 *attCs*, averaging 14.3 per genome). Cassette composition is highly variable even among strains of the same pathovar, suggesting that, in *X. campestris*, integron activity drives rapid diversification within genetically related groups. This robust activity is also evident in the gene content: 216 orthologous cassette-encoded genes were identified, 41.1% of which are singletons (observed in only one instance in this dataset). Despite this extensive variability, a gene encoding a DUF1488 domain-containing protein is present in the cassettes of 81.5% of genomes, while *symE* (a toxin-encoding gene) is found in 56.8% of the cassette arrays. This level of integron retention suggests that active integrons continue to play a key role in the adaptive evolution of *X. campestris*.

*X. arboricola* exhibited a modest preservation of the *intI* gene with 7% of strains retaining a full-length *intI.* In the remaining genomes, *intI* is either deleted or contains an early stop codon. The *X. arboricola* phylogeny divides strains into two clades, clade A which includes pathovars that cause diseases in *Prunus*, *Juglans*, and *Corylus* spp. (pvs. *pruni*, *juglandis*, and *corylina*, respectively), and clade B which comprises many isolates lacking metadata (Figure S3). Among clade A isolates, none possess a full-length *intI.* In contrast, clade B strains that carry *intI*, either full-length or truncated (accounting for 58.8% of clade B isolates), harbor a variant likely acquired via HGT, suggesting the acquisition event dated prior to the diversification of this clade. Strains of clade B also carried more cassettes and a broader array of genes compared to clade A.

*X. translucens* (Figure S4) and *X. euvesicatoria* (Figure S5) emphasize the dynamic nature of integrons in *Xanthomonas.* Both species exhibit high frequencies of full-length *intI* (22.7% and 45.6% respectively) and variability in cassette content is high (44.6% and 50% respectively).

*X. oryzae* includes pathogens of rice (pv. *oryzae* and pv. *oryzicola*), and pathogens of a pervasive weed species that grows along rivers and canals surrounding rice paddies (pv. *leersiae*). However, only X11-5A, a weakly pathogenic strain (Triplett *et al*., 2011) distantly related to the above-mentioned pathovars carries a full-length *intI*. Divergent cassette compositions were observed among the pathovars (Figure S6). This supports the hypothesis that integron activity in *X. oryzae* was ancestrally active, followed by progressive inactivation concomitant with pathovar diversification.

None of the *X. cissicola* (Figure 4) and X. *phaseoli* (Figure S7) isolates retain a full-length, functional *intI*. However, 28.7% and 38% of the genes carried by the cassettes in these species, respectively, appear in one isolate only (singletons). *X. cissicola* comprises isolates commonly known as *X. citri*, an important pathogen able to infect many plants including *Citrus* (pv. *citri*). *X. citri* pv. *citri* includes three recognized pathotypes: A, A*, and AwA. All pv. *citri* genomes analysed here, which include both A and AwA pathotypes, harbor a CALIN element adjacent to *secD* and lack the integrase gene *intI*. All the CALINs in the pv. *citri* contain the same two gene cassettes, which are not present in any other genome analysed in this study, suggesting that the integron platform was inactivated prior to pathotype diversification (Figure 4). A recent genomic study of 95 pv. *citri* strains estimated that the diversification of these pathotypes occurred 1,730 to 5,663 years ago (Patané *et al*., 2019). Given that citrus domestication is thought to have occurred at least 2,000 years ago (Rao *et al*., 2021), the inactivation event likely predates the domestication of the host genus. One of the two cassettes encodes a protein homologous to members of the late embryogenesis abundant (LEA) protein family, which is known to confer protection against water deficit in bacteria (Raga-Carbajal *et al*., 2022). This indicates that the cassette’s function could be oriented toward general environmental stress rather than mediating specific interactions with the plant host.

### Gene cassette abundance is species specific

We investigated whether variation in integron cassette array size, quantified by the number of *attC* sites per genome (Table S1), is primarily shaped by phylogeny or by the ecological context of isolation, categorized by cultivation type (crop, non-crop, crop tree, or ornamental host plants) (Table S4). We used only the species that had more than 10 genomes in our dataset, additionally, we excluded genomes lacking information on the host of isolation, and one strain isolated from mud. We first assessed whether variation in *attC* site counts exhibited a phylogenetic signal. Pagel’s λ test indicated a very strong and highly significant phylogenetic signal (λ=0.99, p < 2.22e-16), indicating that *attC* abundance is strongly structured by evolutionary history. Given this strong phylogenetic structure, we applied a phylogenetic generalized least squares regression to evaluate the effect of cultivation type while accounting for shared ancestry. Isolates from ornamental plants had significantly more *attC* sites (p < 0.001; mean = 12.1), whereas those from tree-crop environments had fewer (p ≈ 0.017; mean = 3.05). No significant differences were observed for crop (mean = 6.01) and non-crop (mean = 9.77) isolates (Figure S9). Overall, these results suggest that while integron array size is strongly shaped by phylogenetic inertia, reflecting higher similarity within rather than between species, ecological factors, such as characteristics of the plant host, may also contribute. The prevalence of a functional integron integrase appears to influence cassette abundance: species with predominantly inactive integrases tend to carry fewer *attC* sites, likely due to gradual cassette loss. However, the evolutionary forces determining why some lineages retain active *intI* while others don’t are unclear.

### Cassette arrays in *Xanthomonas* harbor diverse but poorly characterized gene functions

Prediction of the taxonomic origins of integron *attC*s was performed with a classification tool developed for *attC*s, which uses both sequence and structural homology information (Ghaly *et al*., 2021), to identify matches to known *attC* ‘types’. The method classified all *attC* sites in the *Xanthomonas* strains analysed as originating within the *Xanthomonadales*. Therefore, there is no evidence of long-distance acquisition of cassettes, and the cassettes appear to circulate at least within the *Xanthomonadales* family.

Arrays greatly varied in length across genomes, with a maximum of 46 cassettes in a single genome (*X*. *arboricola_F* GCA_040182365.1) and 33 cassettes in a single array (*X*. *arboricola* GCA_041475745.1 and GCA_041475785.1) (Table S1). Despite the majority of the isolates carrying inactive integron modules, with *intI* deleted or truncated, the cassette arrays appear to have been extremely dynamic before the inactivation of the integron integrase.

Overall, the 706 *Xanthomonas* genomes carried 1004 cassettes arrays, with a total of 4,773 cassettes, within these, Proteinortho (Klemm *et al*., 2023) identified 1,087 different orthologous genes. Of the 1,087 orthologous genes identified by Proteinortho, 476 (43.8%) were present as singletons, unique to one isolate.

Of the 1,087 orthologous genes, only 11.9% could be classified into a known Clusters of Orthologous Groups of proteins (COG) category (Galperin *et al*., 2025) (Table S5). This underrepresentation of known COGs among cassette proteins is well known and has been reported for integrons from a range of different hosts and environments (Rowe-Magnus *et al*., 2003, Mazel, 2006, Boucher *et al*., 2007, Ghaly *et al*., 2024, Rajabal *et al*., 2024). This may be due to sampling bias in databases, with cap tured cassettes potentially being fromuncharacterised environmental organisms (Rodríguez Del Río *et al*., 2024), or may be a result of cassette genes being subject to high mutation rates after capture.

Of the most represented functions, excluding unknown and recombination and repair, 1.47% of the total cassette-encoded proteins were predicted to play a role in transcription, 0.9% in amino acid transport and metabolism, and 0.55% in defence mechanisms. To gain further insight into possible cassette functions, we performed both sequence-based homology searching against eggNOG 5.0 (Huerta-Cepas *et al*., 2019), and protein structural homology searching against the Swiss-Prot database. From this combined approach, putative functions could only be assigned to 6.8% of the proteins. Of these annotated cassette-encoded proteins, 47.7% were classified either as transposase, integrases or insertion sequences (4.8% of the total). The presence of insertion sequences and transposases within gene cassettes (Ghaly *et al*., 2024), and targeting *attC* sites (Tetu & Holmes, 2008, Post & Hall, 2009), has been frequently observed in past integron studies.

Signal peptides were identified in 11.6% of cassette-encoded proteins, using signalP6.0 (Teufel *et al*., 2022). Again, transmembrane and secreted proteins are commonly encoded by gene cassettes in Bacteria (Rowe-Magnus *et al*., 2003, Ghaly *et al*., 2023) and are hypothesized to help facilitate interactions with their broader environment. Additionally, PGPT-Pred (Ashrafi *et al*., 2022) predicted 6.7% of cassette-encoded proteins to have plant growth promoting traits such as biofertilization, plant signal production and stress control. However, the most frequent category, representing 4.9% of total proteins, was associated with competitive exclusion functions. These included antimicrobial resistance and detoxification functions, and toxin-antitoxin systems, which in a plant pathogenic context could contribute to niche adaptation by enhancing survival in a competitive microbial environment. One gene was classified by the EcoFoldDB (Ghaly *et al*., 2025) pipeline as involved in spermidine production, which can play an important role in plant growth and stress (Dunn & Becerra-Rivera, 2023).

DefenseFinder (Tesson *et al*., 2022) identified 20 orthologous genes (1.8%) as genes involved in defences against phages. Twelve different defence systems were detected, these included restriction-modification and abortive infection systems, but also the newly described Kiwa (Zhang *et al*., 2023) and Shedu defence systems (Loeff *et al*., 2025).

Using a database of previously characterized type three secretion system effectors (T3SEs) retrieved from the EuroXanth platform (Costa *et al*., 2024, Biessy *et al*., 2025), we identified two T3SEs: AvrBs3 (a Transcription Activator-Like effector, TALe) in African-like pv. *oryzae* (Schepler-Luu *et al*., 2023), and XopAF2 in three *X. vasicola* isolates. T3SEs are protein secreted via the type III secretion system (T3SS) that suppress or induce plant defences.TALes are a particular type of T3SE, once inside the host cell, they translocate to the nucleus where their unique domain of tandemly arranged 34-amino acid repeats mediates binding to specific promoter elements. Several TALes are known to induce host SWEET sucrose uniporter genes, thereby facilitating sucrose efflux from xylem parenchyma into the apoplasm at the infection sites. TALes may enh ance disease by targeting susceptibility genes, or may trigger a resistance response, and are therefore important for pathogen host range and virulence (Schepler-Luu *et al*., 2023).

Given that the majority of cassette-encoded proteins are of unknown function and often occur as singletons (observed only in one strain), the phenotypic impact of these cassettes remains unclear. It is uncertain whether they confer a selective advantage or are maintained in the arrays through neutral processes. A cassette that is conserved across multiple species is more likely to encode a beneficial function. The most widespread cassette, identified in 13 species, encodes *symE* and the non-coding small RNA *sRNA-Xcc1*. *sRNA-Xcc1* is known to be activated by regulatory elements of the T3SS (Chen *et al*., 2011), suggesting a potential role in plant host colonization. In *E. coli*, *symE* forms part of the SymE-SymR toxin-antitoxin system, where SymE induces nucleoid condensation, disrupts DNA replication and transcription, and causes double-strand DNA breaks (Thompson *et al*., 2022). Regulation in *E. coli* occurs via the cis-encoded small RNA *symR* (Kawano *et al*., 2007), whereas in the integron cassettes, *sRNA-Xcc1* is located upstream of *symE*, suggesting a divergent regulatory mechanism. Notably, unlike most integron-associated genes oriented to be transcribed from the *Pc* promoter (typically located within *intI*), both *sRNA-Xcc1* and *symE* are oriented in the opposite direction and not under *Pc* control.

Excluding transposases, the second most recurrent cassette, shared among 8 species, encoded a protein of unknown function. The third most recurrent, present in 7 species, included two hypothetical proteins and a gene conferring resistance to bleomycin.

Bleomycin resistance genes have been previously identified as enriched in gene cassettes relative to total metagenomes in environmental samples, as shown by both cassette-targeted amplicon sequencing and shotgun metagenomics (Ghaly *et al*., 2023). One of the recurring cassettes is an “empty” cassette which can occur multiple times in the same array. This cassette could have promoter activity as it was previously suggested in other species (Michael & Labbate, 2010, Tansirichaiya *et al*., 2019, Blanco *et al*., 2024).

### Detection of inter-species horizontal transfer of cassette arrays

To assess interspecies cassette gene sharing, we examined pairs of species that most frequently share orthologous genes. As expected, species harbouring a greater number of cassettes tend to share more orthologous genes with others. For example, *X. arboricola* and *X. campestris* shared the highest number of orthologous genes (21), with *X. campestris* sharing the most overall (57 orthologous genes across multiple species).

We then looked for more recent evidence of HGT of cassettes present in multiple species. We used Alfy v1.0.5 (Domazet-Lošo & Haubold, 2011) to identify putative regions subject to HGT. We further selected the regions identified by Alfy 1.0.5 selecting those based on the difference in ANI between the cassette and the genomes harbouring them (Figure S10). With this approach, we detected the movement of 95 cassettes between 38 pairs of species, even between group 1 and 2 *Xanthomonas* (Figure 5). Figure 5 shows the number of cassettes exchanged between species, without normalization for genome or cassette abundance. While this approach does not account for differences in genome diversity within and between species, or for uneven sampling depth (i.e. pandemic lineages are often oversampled and appear less diverse in comparison to commensal isolates), it nevertheless provides a direct view of observed horizontal transfer events. The highest flux of cassettes (10) was between *X. arboricola* and *X*. *euroxanthea,* and between *X. campestris* and *X. hortorum.* However, *X. campestris* is the species with the highest flux of cassettes compared to any other species (36). One of the cassettes that showed evidence of horizontal transfer was that carrying *symE* and *sRNA-Xcc1*.

**Figure 5.**
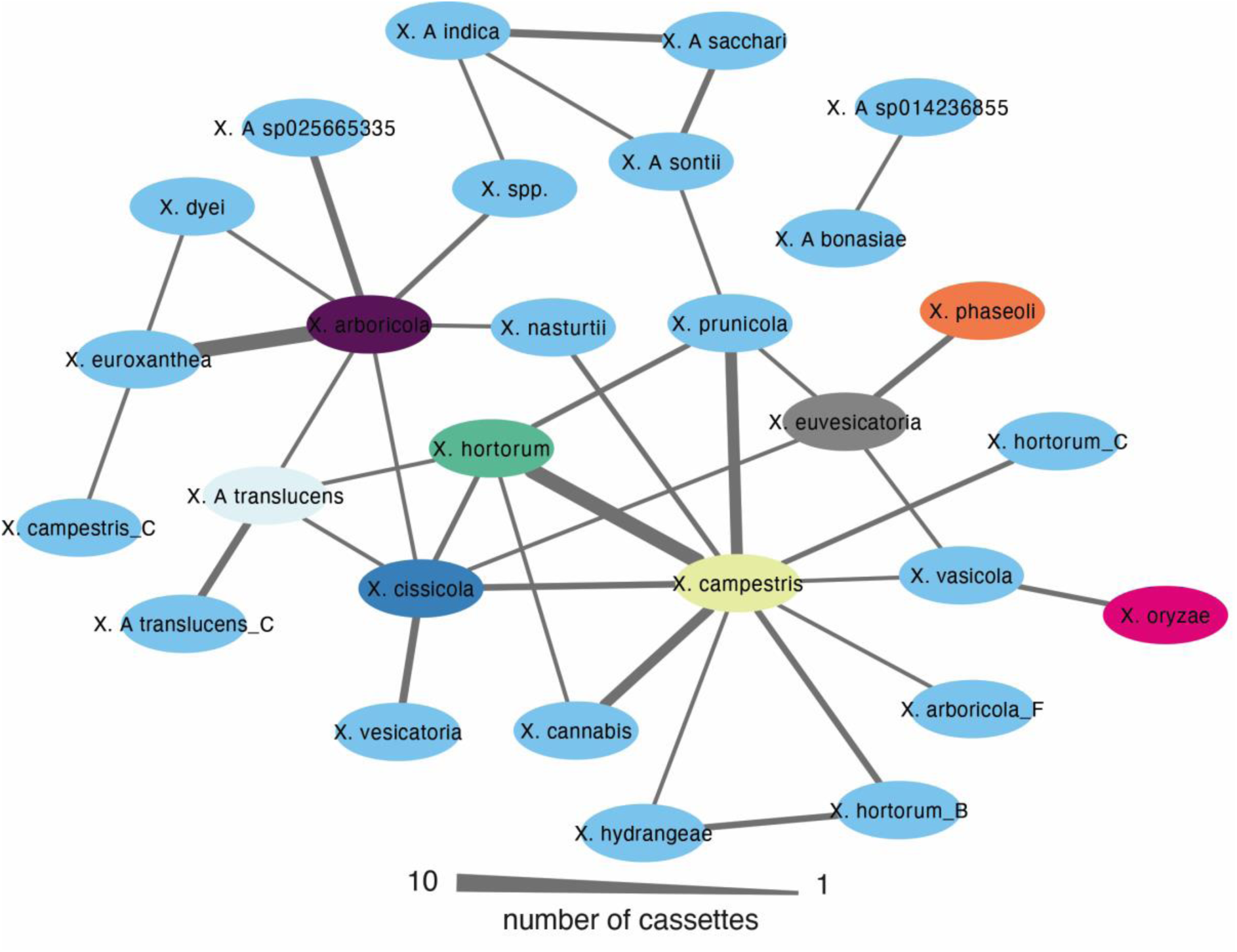
Horizontal transfer of cassettes among *Xanthomonas* species. Cytoscape network illustrating transfer of cassettes between species. Nodes represent *Xanthomonas* species, species containing more than 10 genomes in the dataset are coloured as in Figure 1 (tip labels). Line width between species is proportional to the number of cassettes moved between species.

## CONCLUSIONS

Our comprehensive analysis of 706 complete genomes reveals that the acquisition of the integron platform is likely an ancestral event in the evolution of the *Xanthomonas* genus. The widespread conservation of the *intI* gene downstream of *ilvD* across species, along with *intI* phylogeny, strongly supports the early acquisition of the integron platform. Only in *X. arboricola* clade B, the “original” *intI* appears to have been substituted via HGT with an *intI* from another member of the Xanthomonadaceae family.

Despite the ancestral acquisition, our results show that integron activity, defined by the presence of full-length, potentially functional *intI* genes, and large and diverse cassette arrays, has been progressively lost in many *Xanthomonas* lineages. This inactivation appears to be species-specific and often predates major evolutionary or ecological transitions, such as pathovar diversification, such as in *X. cissicola* and *X. oryzae*. Nevertheless, in species such as *X. campestris*, integrons remain robustly functional and continue to contribute to genomic diversification, as evidenced by the presence of full-length *intIs*, high cassette variability, and the presence of strain-specific gene cassettes in cassette arrays. Selective pressures and ecological niches may favor the retention of an active integron system in this species and not in the others, for reasons not clear at this time. Loss of integron integrase activity in plant-associated pathovars may be favoured when it stabilizes cassette-associated genetic traits that confer a fitness advantage. Once a beneficial cassette configuration is established, inactivation of the integrase can prevent rearrangements or excisions that would disrupt key functions, enabling these lineages to undergo clonal expansion. Integron integrase activity may be also lost in the absence of positive selection to maintain it. Integrons provide the greatest adaptive advantage in dynamic environments with a diverse pool of accessible gene cassettes. Such diversity is potentially diminished in intensive agricultural systems, where pathogens encounter repetitive host-pathogen cycles and microbial communities are less diverse than in wild plant environments (Karasov *et al*., 2018). Under such conditions, the selective pressure to maintain integrase activity might be reduced. In agricultural settings, other types of mobile genetic elements carrying genes already optimized by selection in donor strains may play a more prominent role in driving plant-bacteria evolution (Cesbron *et al*., 2015, Colombi *et al*., 2023, Colombi *et al*., 2024, Jibrin *et al*., 2024, Sparks *et al*., 2025).

The phenotypic impact of the large pool of gene cassettes residing in *Xanthomonas* genomes remains largely unresolved. Nearly half of orthologous genes carried by gene cassettes occurred only once, contributing to strain level genetic variability. These rare genes may represent a reservoir of adaptive potential, but could also be selectively neutral or even transient. Cassettes in *Vibrio* chromosomal integron platforms have been suggested to have been selected to be as neutral as possible (Blanco *et al*., 2024). The frequent occurrence of hypothetical proteins, including highly conserved cassettes ORFs, suggests important functional roles that remain undiscovered. For instance, a cassette carrying the DUF1488 encoding gene is present, a few times in multiple copies, in 16.7% of the genomes across 6 species, sometimes with multiple copies within the same integron array. Among cassette-encoded genes with predicted putative functions, many have predicted functions linked to environmental interaction and thus have the potential to contribute to niche adaptation. For example, the most widespread cassette in our dataset encodes *symE* and the small coding sRNA-Xcc1, known to be activated by T3SS regulatory proteins (Chen *et al*., 2011), which may influence bacterial growth during plant colonization. In *Xanthomonas*, we found a fairly small fraction of cassette genes to be classified as involved defence mechanisms, despite the emerging role of integron platforms in encoding anti-phage genes (Darracq *et al*., 2025, Getz *et al*., 2025, Kieffer *et al*., 2025). However, this may be an underestimate, as work on large sedentary chromosomal integrons in *Vibrio cholerae* showed that 20% of their cassettes with no predicted function encode for anti-phage defence systems, despite these genes not being detected as such by bioinformatic tools (Darracq *et al*., 2025).

Importantly, our analysis revealed evidence of interspecies horizontal transfer of cassettes. For both *X. campestris* and *X. arboricola* there is evidence of recent horizontal transfer of integron-associated cassettes with other *Xanthomonas* species. Interestingly, however, despite their high levels of cassette acquisition, there is little evidence of recent and direct cassette exchange between these two species. Instead, *X. campestris* shows the highest degree of gene sharing with *X. hortorum*, a species more closely related to *X. arboricola* than to *X. campestris*. Conversely, the species with which *X. arboricola* shares the greatest number of cassettes is *X. euroxanthea*, its closest phylogenetic neighbour among the genomes analysed. These patterns suggest that both phylogenetic proximity and ecological overlap, such as co-occurrence in the phyllosphere or endosphere during mixed infections, have contributed to the dynamics of cassette exchange. The shared habitat provides repeated opportunities for interspecies contact and genetic exchange, likely promoting the horizontal dissemination of integron cassettes across the genus.

In summary, our analysis indicates that integron platforms were likely acquired early in the evolution of the *Xanthomonas* genus and have since followed divergent evolutionary trajectories. While many lineages show signs of integron inactivation, others, such as *X. campestris*, retain active systems contributing to genomic diversification. The gene cassettes carried by these integrons, including many with unknown or potentially adaptive functions, add to strain-level variability and may influence niche adaptation. Patterns of horizontal gene transfer suggest that both phylogenetic relatedness and ecological overlap shape cassette exchange across species, highlighting integrons as dynamic elements in the evolutionary landscape of *Xanthomonas*.

## Supporting information

Supplementary Result Section and Supplementary Figures

## FIGURE AND TABLE LEGENDS

**Figure S1 – Phylogeny of *Xanthomonas intI*.** A maximum likelihood phylogenetic tree of all *intI* in *Xanthomonas*. *IntI* from *Vibrio* sp. SCSIO 43136 (locus tag: J4N39_08275) was used as an outgroup. The tree was inferred with FastTree and rooted with *Vibrio* sp. SCSIO 43136, the scale bar indicates substitutions per site. * indicates the *intI*s are not integrated adjacent to *ilvD*.

**Figure S2 – Phylogeny of *ilvD*.** The maximum-likelihood phylogenetic tree of all *ilvD* in *Xanthomonas*. *ilvD* from *Vibrio* sp. SCSIO 43136 was used as an outgroup. The tree was inferred with FastTree and rooted at midpoint, the scale bar indicates substitutions per site.

**Figure S3 – Integrons in *X. arboricola*.** The phylogeny of *X. arboricola* was built using Realphy, using GCA_000972745.1 as a reference genome. The tree was rooted with GCA_001908755.1 (*X. hortorum*) then the tip was removed from the tree. Tip labels show pathovars, which were assigned via literature search (Table S3). The red triangle indicates the probable acquisition of the *intI* variant. The scale bar indicates substitutions per site. A) Indicates the presence of a complete integron or *attC* sites in the *ilvD* locus, B) indicates the presence of an *intI* lacking cassettes (In0), C) indicates the presence of a CALINs (cluster of *attC* sites) not in the ilvD locus, D) indicates whether IntI is predicted to be functional (full length) (Table S3). E) indicates the number of *attC* sites (number of cassettes) in the genome. Panel F) represents the distribution of orthologous genes among the all cassettes carried by the corresponding isolate inferred with Proteinortho. Orthologous genes are given in decreasing order based on the number of strains within the species that carry them.

**Figure S4 – Integrons in *X. translucens*.** The phylogeny of *X. translucens* was built using Realphy, using GCA_001021935.1 as reference genome. The tree was rooted with GCA_000007145.1 (*X. campestris*) then the tip removed from the tree. Tip labels show pathovars, which were assigned via literature search (Table S3). The scale bar indicates substitutions per site. A) Indicates the presence of a complete integron or *attC* sites in the *ilvD* locus, B) inidicates the presence of an *intI* lacking cassettes (In0), C) indicate the presence of a CALINs (cluster of *attC* sites) not in the *ilvD* locus, D) indicates whether IntI is predicted to be functional (full length) (Table S3). E) indicates the number of *attC* sites (number of cassettes) in the genome. Panel F) represents the distribution of orthologous genes among the all the cassettes carried by the corresponding isolate interfered with Proteinortho. Orthologous genes are ordered in decreasing order based on the number of strains within the species that carry them.

**Figure S5 – Integrons in *X. euvesicatoria*.** The phylogeny of *X. euvesicatoria* was built using Realphy, using GCA_000009165.1 as reference genome. The tree was rooted with GCA_002759095.2 (*X. phaseoli*) then the tip removed from the tree. The scale bar indicates substitutions per site. A) Indicates the presence of a complete integron or *attC* sites in the *ilvD* locus, B) inidicates the presence of an *intI* lacking cassettes (In0), C) indicate the presence of a CALINs (cluster of *attC* sites) not in the *ilvD* locus, D) indicates whether IntI is predicted to be functional (full length) (Table S3). E) indicates the number of *attC* sites (number of cassettes) in the genome. Panel F) represents the distribution of orthologous genes among the all the cassettes carried by the corresponding isolate interfered with Proteinortho. Orthologous genes are ordered in decreasing order based on the number of strains within the species that carry them.

**Figure S6 – Integrons in *X. oryzae*.** The phylogeny of *X. oryzae* was built using Realphy, using GCA_000007385.1 as reference genome. The tree was rooted with GCA_002759095.2 (*X. phaseoli*) then the tip removed from the tree. Tip labels show pathovars, which were assigned via literature search (Table S3). The scale bar indicates substitutions per site. A) Indicates the presence of a complete integron or *attC* sites in the *ilvD* locus, B) inidicates the presence of an *intI* lacking cassettes (In0), C) indicate the presence of a CALINs (cluster of *attC* sites) not in the *ilvD* locus, D) indicates whether IntI is predicted to be functional (full length). E) indicates the number of *attC* sites (number of cassettes) in the genome (Table S3). Panel F) represents the distribution of orthologous genes among the all the cassettes carried by the corresponding isolate interfered with Proteinortho. Orthologous genes are ordered in decreasing order based on the number of strains within the species that carry them.

**Figure S7 – Integrons in *X. phaseoli*.** The phylogeny of *X. phaseoli* was built using Realphy, using GCA_008639345.1 as reference genome. The tree was rooted with GCA_000009165.1 (*X. euvesicatoria*) then the tip removed from the tree. Tip labels show pathovars, which were assigned via literature search (Table S3). The scale bar indicates substitutions per site. A) Indicates the presence of a complete integron or *attC* sites in the *ilvD* locus, B) inidicates the presence of *intI* lacking cassettes (In0), C) indicate the presence of a CALINs (cluster of *attC* sites) not in the *ilvD* locus, D) indicates whether IntI is predicted to be functional (full length) (Table S3). E) indicates the number of *attC* sites (number of cassettes) in the genome. Panel F) represents the distribution of orthologous genes among the all the cassettes carried by the corresponding isolate interfered with Proteinortho. Orthologous genes are ordered in decreasing order based on the number of strains within the species that carry them.

**Figure S8 – Integrons in *X. hortorum*.** The phylogeny of *X. hortorum* was built using Realphy, using GCA_002285515.1 as reference genome. The tree was rooted with GCA_905367715.1 (*X. arboricola*) then the tip removed from the tree. Tip labels show pathovars, which were assigned via literature search (Table S3). The scale bar indicates substitutions per site. A) Indicates the presence of a complete integron or *attC* sites in the *ilvD* locus, B) indicates the presence of an *intI* lacking cassettes (In0), C) indicate the presence of a CALINs (cluster of *attC* sites) not in the *ilvD* locus, D) indicates whether IntI is predicted to be functional (full length). E) indicates the number of *attC* sites (number of cassettes) in the genome (Table S3). Panel F) represents the distribution of orthologous genes among the all the cassettes carried by the corresponding isolate interfered with Proteinortho. Orthologous genes are ordered in decreasing order based on the number of strains within the species that carry them.

**Figure S9 – Integron gene cassette counts in *Xanthomonas*.** Boxplot of the number of *attC* sites per genome grouped by plant of isolation and by species (only species with more than 10 genomes are shown).

**Figure S10 – Delta ANI values for selected events of interspecies transfer of cassettes.** A) Dot plot and B) density plot of the cassette-to-genome ANI differences (delta ANI). In A), black dots represent individual data points, while in B), the curve shows the density estimate of the distribution.

**Table S1 – *Xanthomonas* genomes used in this study with integrons and number of cassettes identified.**

**Table S2 – Coordinates of *intI* identified in *Xanthomonas*.**

**Table S3 – Selected *Xanthomonas* genomes with integrons and number of cassettes identified, pathovar information, and predicted *intI* functionality.**

**Table S4 – Metadata associated with Selected *Xanthomonas* genomes.**

**Table S5 – Identified COG categories of the 1,087 orthologous genes in non-redundant cassettes.**

## REFERENCES

Almagro Armenteros JJ, Tsirigos KD, Sønderby CK, Petersen TN, Winther O, Brunak S, von Heijne G & Nielsen H (2019) SignalP 5.0 improves signal peptide predictions using deep neural networks. Nature biotechnology 37: 420–423.

Ansari MA & Didelot X (2014) Inference of the properties of the recombination process from whole bacterial genomes. Genetics 196: 253–265.

Ashrafi S, Kuzmanović N, Patz S, Lohwasser U, Bunk B, Spröer C, Lorenz M, Elhady A, Frühling A & Neumann-Schaal M (2022) Two new Rhizobiales species isolated from root nodules of common sainfoin (Onobrychis viciifolia) show different plant colonization strategies. Microbiology Spectrum 10: e01099–01022.

Barionovi D & Scortichini M (2006) Assessment of integron gene cassette arrays in strains of *Xanthomonas fragariae* and *X. arboricola* pvs. *fragariae* and *pruni*. Journal of Plant Pathology 279–284.

Barionovi D & Scortichini M (2008) Integron variability in *Xanthomonas arboricola* pv. *juglandis* and *Xanthomonas arboricola* pv. *pruni* strains. FEMS microbiology letters 288: 19–24.

Bertels F, Silander OK, Pachkov M, Rainey PB & Van Nimwegen E (2014) Automated reconstruction of whole-genome phylogenies from short-sequence reads. Molecular biology and evolution 31: 1077–1088.

Biessy A, Cadieux M, Ciotola M, McDuff F, Soufiane B, Laforest M & Filion M (2025) Characterization of a plant-pathogenic T3SS-lacking *Xanthomonas* strain isolated from common ragweed. Plant Pathology 74: 308–319.

Blanco P, Hipólito A, García-Pastor L, Trigo da Roza F, Toribio-Celestino L, Ortega AC, Vergara E, San Millán Á & Escudero JA (2024) Identification of promoter activity in gene-less cassettes from *Vibrionaceae* superintegrons. Nucleic acids research 52: 2961–2976.

Boucher Y, Labbate M, Koenig JE & Stokes H (2007) Integrons: mobilizable platforms that promote genetic diversity in bacteria. Trends in microbiology 15: 301–309.

Buchfink B, Xie C & Huson DH (2015) Fast and sensitive protein alignment using DIAMOND. Nature methods 12: 59–60.

Cambray G, Sanchez-Alberola N, Campoy S, Guerin É, Da Re S, González-Zorn B, Ploy M-C, Barbé J, Mazel D & Erill I (2011) Prevalence of SOS-mediated control of integron integrase expression as an adaptive trait of chromosomal and mobile integrons. Mobile DNA 2: 6.

Cantalapiedra CP, Hernández-Plaza A, Letunic I, Bork P & Huerta-Cepas J (2021) eggNOG-mapper v2: functional annotation, orthology assignments, and domain prediction at the metagenomic scale. Molecular biology and evolution 38: 5825–5829.

Cesbron S, Briand M, Essakhi S, Gironde S, Boureau T, Manceau C, Fischer-Le Saux M & Jacques M-A (2015) Comparative genomics of pathogenic and nonpathogenic strains of *Xanthomonas arboricola* unveil molecular and evolutionary events linked to pathoadaptation. Frontiers in Plant Science 6: 1126.

Chaumeil P-A, Mussig AJ, Hugenholtz P & Parks DH (2022) GTDB-Tk v2: memory friendly classification with the genome taxonomy database. Bioinformatics 38: 5315–5316.

Chen X-L, Tang D-J, Jiang R-P, He Y-Q, Jiang B-L, Lu G-T & Tang J-L (2011) *sRNA-Xcc1*, an integron-encoded transposon-and plasmid-transferred trans-acting sRNA, is under the positive control of the key virulence regulators HrpG and HrpX of *Xanthomonas campestris* pathovar *campestris*. RNA biology 8: 947–953.

Colombi E, Hill Y, Lines R, Sullivan JT, Kohlmeier MG, Christophersen CT, Ronson CW, Terpolilli JJ & Ramsay JP (2023) Population genomics of Australian indigenous *Mesorhizobium* reveals diverse nonsymbiotic genospecies capable of nitrogen-fixing symbioses following horizontal gene transfer. Microbial genomics 9: 000918.

Colombi E, Bertels F, Doulcier G, McConnell E, Pichugina T, Sohn KH, Straub C, McCann HC & Rainey PB (2024) Rapid dissemination of host metabolism–manipulating genes via integrative and conjugative elements. Proceedings of the National Academy of Sciences 121: e2309263121.

Costa J, Pothier JF, Bosis E, Boch J, Kölliker R & Koebnik R (2024) A community-curated DokuWiki resource on diagnostics, diversity, pathogenicity, and genetic control of xanthomonads. Molecular Plant-Microbe Interactions 37: 347–353.

Cury J, Jové T, Touchon M, Néron B & Rocha EP (2016) Identification and analysis of integrons and cassette arrays in bacterial genomes. Nucleic acids research 44: 4539–4550.

Darracq B, Littner E, Brunie M, Bos J, Kaminski PA, Depardieu F, Slesak W, Debatisse K, Touchon M & Bernheim A (2025) Sedentary chromosomal integrons as biobanks of bacterial antiphage defense systems. Science 388: eads0768.

Domazet-Lošo M & Haubold B (2011) Alignment-free detection of local similarity among viral and bacterial genomes. Bioinformatics 27: 1466–1472.

Dunn MF & Becerra-Rivera VA (2023) The biosynthesis and functions of polyamines in the interaction of plant growth-promoting rhizobacteria with plants. Plants 12: 2671.

Escudero JA, Loot C & Mazel D (2018) Integrons as adaptive devices. Molecular mechanisms of microbial evolution 199–239.

Essakhi S, Cesbron S, Fischer-Le Saux M, Bonneau S, Jacques M-A & Manceau C (2015) Phylogenetic and variable-number tandem-repeat analyses identify nonpathogenic Xanthomonas arboricola lineages lacking the canonical type III secretion system. Applied and Environmental Microbiology 81: 5395–5410.

Galperin MY, Vera Alvarez R, Karamycheva S, Makarova KS, Wolf YI, Landsman D & Koonin EV (2025) COG database update 2024. Nucleic Acids Research 53: D356–D363.

Garita-Cambronero J, Palacio-Bielsa A, López MM & Cubero J (2017) Pan-genomic analysis permits differentiation of virulent and non-virulent strains of Xanthomonas arboricola that cohabit Prunus spp. and elucidate bacterial virulence factors. Frontiers in microbiology 8: 573.

Getz LJ, Fairburn SR, Vivian Liu Y, Qian AL & Maxwell KL (2025) Integrons are anti-phage defence libraries in *Vibrio parahaemolyticus*. Nature Microbiology 1–10.

Ghaly TM, Tetu SG & Gillings MR (2021) Predicting the taxonomic and environmental sources of integron gene cassettes using structural and sequence homology of attC sites. Communications biology 4: 946.

Ghaly TM, Geoghegan JL, Alroy J & Gillings MR (2019) High diversity and rapid spatial turnover of integron gene cassettes in soil. Environmental Microbiology 21: 1567–1574.

Ghaly TM, Geoghegan JL, Tetu SG & Gillings MR (2020) The peril and promise of integrons: beyond antibiotic resistance. Trends in microbiology 28: 455–464.

Ghaly TM, Gillings MR, Rajabal V, Paulsen IT & Tetu SG (2024) Horizontal gene transfer in plant microbiomes: integrons as hotspots for cross-species gene exchange. Frontiers in Microbiology 15: 1338026.

Ghaly TM, Rajabal V, Russel D, Colombi E & Tetu SG (2025) EcoFoldDB: Protein structure-guided functional profiling of ecologically relevant microbial traits at the metagenome scale. bioRxiv 2025.2004.2002.646905.

Ghaly TM, Tetu SG, Penesyan A, Qi Q, Rajabal V & Gillings MR (2022) Discovery of integrons in Archaea: Platforms for cross-domain gene transfer. Science Advances 8: eabq6376.

Ghaly TM, Penesyan A, Pritchard A, Qi Q, Rajabal V, Tetu SG & Gillings MR (2022) Methods for the targeted sequencing and analysis of integrons and their gene cassettes from complex microbial communities. Microbial genomics 8: 000788.

Ghaly TM, Rajabal V, Penesyan A, Coleman NV, Paulsen IT, Gillings MR & Tetu SG (2023) Functional enrichment of integrons: Facilitators of antimicrobial resistance and niche adaptation. Iscience 26.

Gillings M, Boucher Y, Labbate M, Holmes A, Krishnan S, Holley M & Stokes H (2008) The evolution of class 1 integrons and the rise of antibiotic resistance. Journal of bacteriology 190: 5095–5100.

Gillings MR (2014) Integrons: past, present, and future. Microbiology and molecular biology reviews 78: 257–277.

Gillings MR, Holley MP, Stokes H & Holmes AJ (2005) Integrons in *Xanthomonas*: a source of species genome diversity. Proceedings of the National Academy of Sciences 102: 4419–4424.

Guerin É, Cambray G, Sanchez-Alberola N, Campoy S, Erill I, Da Re S, Gonzalez-Zorn B, Barbé J, Ploy M-C & Mazel D (2009) The SOS response controls integron recombination. Science 324: 1034–1034.

Hall RM & Collis CM (1995) Mobile gene cassettes and integrons: capture and spread of genes by site-specific recombination. Molecular microbiology 15: 593–600.

Heinzinger M, Weissenow K, Sanchez JG, Henkel A, Mirdita M, Steinegger M & Rost B (2024) Bilingual language model for protein sequence and structure. NAR Genomics and Bioinformatics 6: lqae150.

Hipólito A, García-Pastor L, Vergara E, Jové T & Escudero JA (2023) Profile and resistance levels of 136 integron resistance genes. npj Antimicrobials and Resistance 1: 13.

Huerta-Cepas J, Szklarczyk D, Heller D, Hernández-Plaza A, Forslund SK, Cook H, Mende DR, Letunic I, Rattei T & Jensen LJ (2019) eggNOG 5.0: a hierarchical, functionally and phylogenetically annotated orthology resource based on 5090 organisms and 2502 viruses. Nucleic acids research 47: D309–D314.

Jain C, Rodriguez-R LM, Phillippy AM, Konstantinidis KT & Aluru S (2018) High throughput ANI analysis of 90K prokaryotic genomes reveals clear species boundaries. Nature communications 9: 5114.

Jibrin MO, Sharma A, Mavian CN, Timilsina S, Kaur A, Iruegas-Bocardo F, Potnis N, Minsavage GV, Coutinho TA & Creswell TC (2024) Phylodynamic insights into global emergence and diversification of the tomato pathogen *Xanthomonas hortorum* pv. *gardneri*. Molecular Plant-Microbe Interactions 37: 712–720.

Karasov TL, Almario J, Friedemann C, et al. (2018) *Arabidopsis thaliana* and *Pseudomonas* pathogens exhibit stable associations over evolutionary timescales. Cell host & microbe 24: 168–179. e164.

Kawano M, Aravind á & Storz G (2007) An antisense RNA controls synthesis of an SOS-induced toxin evolved from an antitoxin. Molecular microbiology 64: 738–754.

Kieffer N, Hipólito A, Ortiz-Miravalles L, Blanco P, Delobelle T, Vizuete P, Ojeda FM, Jové T, Jurenas D & García-Quintanilla M (2025) Mobile integrons encode phage defense systems. Science 388: eads0915.

Klemm P, Stadler PF & Lechner M (2023) Proteinortho6: pseudo-reciprocal best alignment heuristic for graph-based detection of (co-) orthologs. Frontiers in bioinformatics 3: 1322477.

Langmead B & Salzberg SL (2012) Fast gapped-read alignment with Bowtie 2. Nature methods 9: 357–359.

Lemoine F & Gascuel O (2021) Gotree/Goalign: toolkit and Go API to facilitate the development of phylogenetic workflows. NAR Genomics and Bioinformatics 3: lqab075.

Loeff L, Walter A, Rosalen GT & Jinek M (2025) DNA end sensing and cleavage by the Shedu anti-phage defense system. Cell 188: 721–733. e717.

Loot C, Millot GA, Richard E, Littner E, Vit C, Lemoine F, Néron B, Cury J, Darracq B & Niault T (2024) Integron cassettes integrate into bacterial genomes via widespread non-classical attG sites. Nature Microbiology 9: 228–240.

Löytynoja A (2013) Phylogeny-aware alignment with PRANK. Multiple sequence alignment methods,p.^pp. 155–170. Springer.

Mazel D (2006) Integrons: agents of bacterial evolution. Nature Reviews Microbiology 4: 608–620.

Mazel D, Dychinco B, Webb VA & Davies J (1998) A distinctive class of integron in the *Vibrio cholerae* genome. Science 280: 605–608.

Michael CA & Labbate M (2010) Gene cassette transcription in a large integron-associated array. BMC genetics 11: 1–13.

Néron B, Littner E, Haudiquet M, Perrin A, Cury J & Rocha EP (2022) IntegronFinder 2.0: identification and analysis of integrons across bacteria, with a focus on antibiotic resistance in Klebsiella. Microorganisms 10: 700.

Orme D, Freckleton R, Thomas G, Petzoldt T, Fritz S, Isaac N & Pearse W (2013) The caper package: comparative analysis of phylogenetics and evolution in R. R package version 5: 1–36.

Paradis E & Schliep K (2019) ape 5.0: an environment for modern phylogenetics and evolutionary analyses in R. Bioinformatics 35: 526–528.

Parks DH, Chuvochina M, Rinke C, Mussig AJ, Chaumeil P-A & Hugenholtz P (2022) GTDB: an ongoing census of bacterial and archaeal diversity through a phylogenetically consistent, rank normalized and complete genome-based taxonomy. Nucleic acids research 50: D785–D794.

Patané JS, Martins J, Rangel LT, Belasque J, Digiampietri LA, Facincani AP, Ferreira RM, Jaciani FJ, Zhang Y & Varani AM (2019) Origin and diversification of *Xanthomonas citri* subsp. *citri* pathotypes revealed by inclusive phylogenomic, dating, and biogeographic analyses. BMC genomics 20: 1–23.

Pena MM, Bhandari R, Bowers RM, Weis K, Newberry E, Wagner N, Pupko T, Jones JB, Woyke T & Vinatzer BA (2024) Genetic and functional diversity help explain pathogenic, weakly pathogenic, and commensal lifestyles in the genus Xanthomonas. Genome Biology and Evolution 16: evae074.

Pereira BM, Österlund T, Eriksson KM, Backhaus T, Axelson-Fisk M & Kristiansson E (2020) A comprehensive survey of integron-associated genes present in metagenomes. BMC genomics 21: 1–14.

Pinheiro J, Bates D, DebRoy S, Sarkar D, Heisterkamp S, Van Willigen B & Maintainer R (2017) Package ‘nlme’. Linear and nonlinear mixed effects models, version 3: 274.

Post V & Hall RM (2009) Insertion sequences in the IS 1111 family that target the attC recombination sites of integron-associated gene cassettes. FEMS microbiology letters 290: 182–187.

Price MN, Dehal PS & Arkin AP (2010) FastTree 2–approximately maximum-likelihood trees for large alignments. PloS one 5: e9490.

Qi Q, Ghaly TM, Rajabal V, Russell DH, Gillings MR & Tetu SG (2024) Vegetable phylloplane microbiomes harbour class 1 integrons in novel bacterial hosts and drive the spread of chlorite resistance. Science of the Total Environment 954: 176348.

Raga-Carbajal E, Espin G, Ayala M, Rodríguez-Salazar J & Pardo-López L (2022) Evaluation of a bacterial group 1 LEA protein as an enzyme protectant from stress-induced inactivation. Applied Microbiology and Biotechnology 106: 5551–5562.

Rajabal V, Ghaly TM, Egidi E, Ke M, Penesyan A, Qi Q, Gillings MR & Tetu SG (2024) Exploring the role of mobile genetic elements in shaping plant–bacterial interactions for sustainable agriculture and ecosystem health. *Plants, People*, Planet 6: 408–420.

Rao MJ, Zuo H & Xu Q (2021) Genomic insights into citrus domestication and its important agronomic traits. Plant Communications 2.

Rodríguez Del Río Á, Giner-Lamia J, Cantalapiedra CP, Botas J, Deng Z, Hernández-Plaza A, Munar-Palmer M, Santamaría-Hernando S, Rodríguez-Herva JJ & Ruscheweyh H-J (2024) Functional and evolutionary significance of unknown genes from uncultivated taxa. Nature 626: 377–384.

Rowe-Magnus DA & Mazel D (2001) Integrons: natural tools for bacterial genome evolution. Current opinion in microbiology 4: 565–569.

Rowe-Magnus DA, Guerout A-M, Biskri L, Bouige P & Mazel D (2003) Comparative analysis of superintegrons: engineering extensive genetic diversity in the Vibrionaceae. Genome research 13: 428–442.

Sapkota S, Mergoum M & Liu Z (2020) The translucens group of Xanthomonas translucens: Complicated and important pathogens causing bacterial leaf streak on cereals. Molecular plant pathology 21: 291–302.

Schepler-Luu V, Sciallano C, Stiebner M, Ji C, Boulard G, Diallo A, Auguy F, Char SN, Arra Y & Schenstnyi K (2023) Genome editing of an African elite rice variety confers resistance against endemic and emerging Xanthomonas oryzae pv. oryzae strains. Elife 12: e84864.

Schwengers O, Jelonek L, Dieckmann MA, Beyvers S, Blom J & Goesmann A (2021) Bakta: rapid and standardized annotation of bacterial genomes via alignment-free sequence identification. Microbial genomics 7: 000685.

Shannon P, Markiel A, Ozier O, Baliga NS, Wang JT, Ramage D, Amin N, Schwikowski B & Ideker T (2003) Cytoscape: a software environment for integrated models of biomolecular interaction networks. Genome research 13: 2498–2504.

Sparks AH, Adorada DL, Colombi E, Kelly LA, Young A, Knight NL & Vaghefi N (2025) Clonal expansion from standing genetic variation underpins the evolution of *Curtobacterium flaccumfaciens* pv. *flaccumfaciens* in Australia. Phytopathology.

Stokes Ht & Hall R (1989) A novel family of potentially mobile DNA elements encoding site-specific gene-integration functions: integrons. Molecular microbiology 3: 1669–1683.

Strugeon E, Tilloy V, Ploy M-C & Da Re S (2016) The stringent response promotes antibiotic resistance dissemination by regulating integron integrase expression in biofilms. MBio 7: 10.1128/mbio.00868-00816.

Tansirichaiya S, Mullany P & Roberts AP (2019) Promoter activity of ORF-less gene cassettes isolated from the oral metagenome. Scientific Reports 9: 8388.

Tesson F, Hervé A, Mordret E, Touchon M, d’Humières C, Cury J & Bernheim A (2022) Systematic and quantitative view of the antiviral arsenal of prokaryotes. Nature communications 13: 2561.

Tetu SG & Holmes AJ (2008) A family of insertion sequences that impacts integrons by specific targeting of gene cassette recombination sites, the IS 1111-attC group. Journal of bacteriology 190: 4959–4970.

Teufel F, Almagro Armenteros JJ, Johansen AR, Gíslason MH, Pihl SI, Tsirigos KD, Winther O, Brunak S, von Heijne G & Nielsen H (2022) SignalP 6.0 predicts all five types of signal peptides using protein language models. Nature biotechnology 40: 1023–1025.

Thompson MK, Nocedal I, Culviner PH, Zhang T, Gozzi KR & Laub MT (2022) *Escherichia coli* SymE is a DNA-binding protein that can condense the nucleoid. Molecular microbiology 117: 851–870.

Timilsina S, Potnis N, Newberry EA, Liyanapathiranage P, Iruegas-Bocardo F, White FF, Goss EM & Jones JB (2020) Xanthomonas diversity, virulence and plant–pathogen interactions. Nature Reviews Microbiology 18: 415–427.

Triplett LR, Hamilton J, Buell C, Tisserat N, Verdier V, Zink F & Leach J (2011) Genomic analysis of Xanthomonas oryzae isolates from rice grown in the United States reveals substantial divergence from known X. oryzae pathovars. Applied and Environmental Microbiology 77: 3930–3937.

Van Kempen M, Kim SS, Tumescheit C, Mirdita M, Lee J, Gilchrist CL, Söding J & Steinegger M (2024) Fast and accurate protein structure search with Foldseek. Nature biotechnology 42: 243–246.

Vauterin L, Yang P, Alvarez A, Takikawa Y, Roth DA, Vidaver AK, Stall RE, Kersters K & Swings J (1996) Identification of non-pathogenic Xanthomonas strains associated with plants. Systematic and applied microbiology 19: 96–105.

Xu S, Dai Z, Guo P, Fu X, Liu S, Zhou L, Tang W, Feng T, Chen M & Zhan L (2021) ggtreeExtra: compact visualization of richly annotated phylogenetic data. Molecular biology and evolution 38: 4039–4042.

Yu G, Smith DK, Zhu H, Guan Y & Lam TTY (2017) ggtree: an R package for visualization and annotation of phylogenetic trees with their covariates and other associated data. Methods in Ecology and Evolution 8: 28–36.

Zhang Z, Todeschini TC, Wu Y, Kogay R, Naji A, Rodriguez JC, Mondi R, Kaganovich D, Taylor DW & Bravo JP (2023) Kiwa is a bacterial membrane-embedded defence supercomplex activated by phage-induced membrane changes. bioRxiv 2023.2002.2026.530102.

