## Supplementary Result Section and Supplementary Figures for "Adaptative and ancient co-evolution of integrons with *Xanthomonas* genomes"

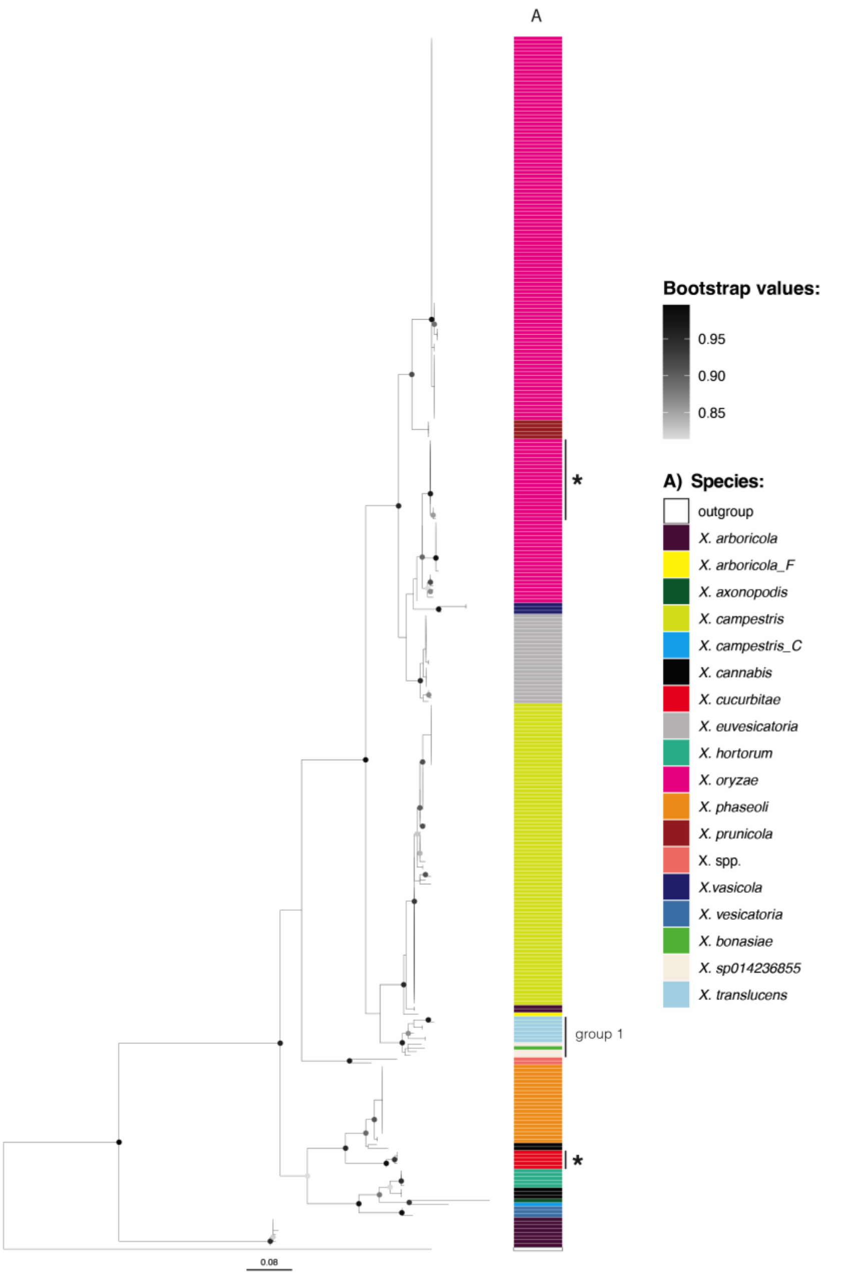


**Figure S1 – Phylogeny of *Xanthomonas intI*.** A maximum likelihood phylogenetic tree of all *intI* in *Xanthomonas*. *IntI* from *Vibrio* sp. SCSIO 43136 (locus tag: J4N39_08275) was used as an outgroup. The tree was inferred with FastTree and rooted with *Vibrio* sp. SCSIO 43136, the scale bar indicates substitutions per site. * indicates the *intI*s are not integrated adjacent to *ilvD*.

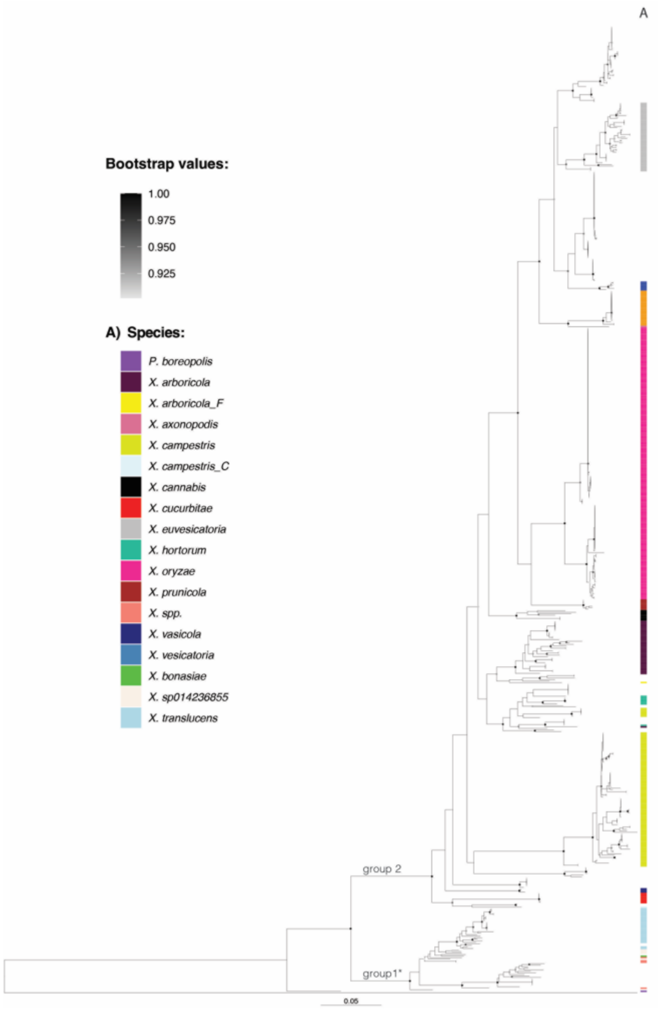


**Figure S2 – Phylogeny of *ilvD*.** The maximum-likelihood phylogenetic tree of all *ilvD* in *Xanthomonas*. *ilvD* from *Vibrio* sp. SCSIO 43136 was used as an outgroup. The tree was inferred with FastTree and rooted at midpoint, the scale bar indicates substitutions per site.

### Integrons in *X. oryzae*, *X. cissicola*, *X. campestris*, *X. euvesicatoria*, *X. translucens*, *X. arboricola*, *X. phaseoli*, and *X. hortorum.*

To investigate integron integrase activity and the complement of integron gene cassettes in *Xanthomonas* genomes, we focused on species which had more than 10 genomes in our dataset: *X. campestris* (n=111) (Figure 3), *X. cissicola* (n= 124) (Figure 4), *X. arboricola* (n=41) (Figure S3), *X. translucens* (n=46) (Figure S4), *X. euvesicatoria* (n=46) (Figure S5), *X. oryzae* (n=161) (Figure S6), *X. phaseoli* (n=23) (Figure S7), *X. hortorum* (n=12) (Figure S8). A detailed description of the integrons in these eight species is reported in the Supplementary Results Section.

#### *X. campestris*

*X. campestris* comprises multiple pathovars that cause distinct diseases on different plant hosts. The pv. *campestris*, the most agriculturally important of these pathovars, is a vascular pathogen causing black rot on *Brassica oleracea* (cabbage and cauliflower), the pv. *raphani* is a nonvascular pathogen causing leaf spot on brassicaceous and solanaceous crops, and the pv. *incanae* is a vascular pathogen of ornamental crucifers (Dubrow *et al.*, 2022).

In comparison to the other *Xanthomonas* species, *X. campestris* maintains a functional integron platform, with 85.6% of the genomes carrying a full-length *intI*, and with all the genomes having from 3 to 29 *attC* sites (average of 14.3 *attCs* per genome).

Integron activity leaves a clear signature in the gene content of the cassettes: out of 216 orthologous genes identified (18 of which are transposases or insertion sequences), the 41.1% of those (91 genes, including 5 transposases) are present in one isolate only (Figure 3).

Cassette composition is highly variable even among strains of the same pathovar, suggesting that, in *X. campestris*, integron activity drives rapid diversification within genetically related groups. Despite this extensive variability, a gene encoding a DUF1488 domain-containing protein is present in the cassettes of 81.5% of genomes, while *symE* is found in 56.8% of the cassette arrays.

#### *X. cissicola*

Although GTDB-Tk classifies this species as *X. cissicola*, it comprises isolates commonly known as *X. citri*, an important pathogen able to infect many plants including *Citrus* (pv. *citri*), legumes such as soybean (pv. *glycines*) and common bean (pv. *fuscans*) (Bansal *et al.*, 2017), and cotton (pv. *malvacearum*) (Delannoy *et al.*, 2005).

The pv. *glycines*, pv. *aurantifolii* lineages XauB and XauC (Fonseca *et al.*, 2019), and one pv. *vigna* strain, which represent 13.7% of the genomes (n=17), have no sign of integrons (no *intI* and no *attC*). *attC* sites are detected in 86.3% of the *X. cissicola* genomes, either in the *ilvD* locus and as CALINs across other three loci. Genomes can carry up to 12 *attCs* with an average of 4 *attC* per genome. In total, the cassettes harbour 87 different orthologous genes, 15 of which are transposases or insertion sequences (ISs), 28.7% of these orthologous genes separately appear in one cassette only (singletons) (25 genes, 4 of which are transposases or ISs) (Figure 4).

*X. citri* pv. *citri* comprises three recognized pathotypes: A, A*, and AwA. A recent genomic study of 95 pv. *citri* strains estimated that the diversification of these pathotypes occurred approximately 1,700 to 5,700 years ago (Patané *et al.*, 2019). All pv. *citri* genomes analysed here, which include both A and AwA pathotypes, harbor a CALIN element adjacent to the *secD* gene and lack the integrase gene *intI*. All the CALINs in the pv. *citri* contain two conserved gene cassettes suggesting that the integron platform was already inactivated prior to pathotype diversification. Given that citrus domestication is thought to have occurred at least 2,000 years ago (Rao *et al.*, 2021), the inactivation event likely predates the domestication of the host plant. One of the two cassettes encodes a protein homologous to members of the late embryogenesis abundant (LEA) protein family, which are known to confer protection against water deficit in bacteria(Raga-Carbajal *et al.*, 2022). This indicates that the cassette's function may be oriented toward general environmental stress rather than mediating specific interactions with the plant host. In contrast to *pv. citri*, which exhibits no cassette diversity, the *pv. fuscans* lineage GL3 and *pv. punicae* strains display a more variable cassette repertoire despite lacking *intI*. Furthermore, two cassettes encoding  proteins of unknown function are present in multiple pathovars, including pvs. *mangiferaeindicae*, *malvacearum*, *punicae*, and *fuscans* lineage GL2.

***X. arboricola***

The 41 genomes analysed here include *X. arboricola* pathovars that cause diseases in *Prunus*, *Juglans*, and *Corylus* spp. (pvs. *pruni*, *juglandis*, and *corylina*, respectively) (clade A, Figure S3) and many isolates whose metadata information are missing (clade B).

Overall, integron elements (*intIs* and or *attC* sites) are present in 85% of the strains, with only six of the eight pv. *corylina* isolates lacking both *attC* sites and *intI*. Only 7% of the strains harbor a full-length *intI*; in the remaining genomes, *intI* is either deleted or contains an early stop codon. Among clade A isolates, which include pathogenic strains, none possess a full-length *intI*, only two pv. *corylina* strains carry a truncated *intI* that is closely related to other *Xanthomonas intIs*. In contrast, clade B strains that carry *intI*, either full-length or truncated (accounting for 58.8% of clade B isolates), harbor the variant likely acquired via HGT, suggesting the event dated prior to the diversification of this clade. On average, *X. arboricola* genomes contain 10 *attC* sites, but clade B isolates also harbours significantly more cassettes ($p<0.01$) then clade A isolates (17±9 and 5±3 cassettes, respectively). Interestingly, the strain with the highest number of *attCs* (33) carries an *intI* harboring an early stop codon resulting from a frameshift mutation. The integron cassette arrays encode a total of 182 unique orthologous genes, of which 4 are transposases or ISs; nearly 45% of these genes appear as singletons. Overall, clade B strains exhibit a higher cassette count compared to clade A strains. This difference may be attributable to the presence of full-length *intIs* in clade B strains, although considerable cassette variability is observed even among isolates with no or only truncated *intIs*. In clade A, it appears that *intI* functionality was lost after the diversification of the various pathovars; within each pathovar, the composition of cassette arrays may vary, although variation is more pronounced in pv. *juglans* in comparison to pvs. *pruni* and *corylina*.

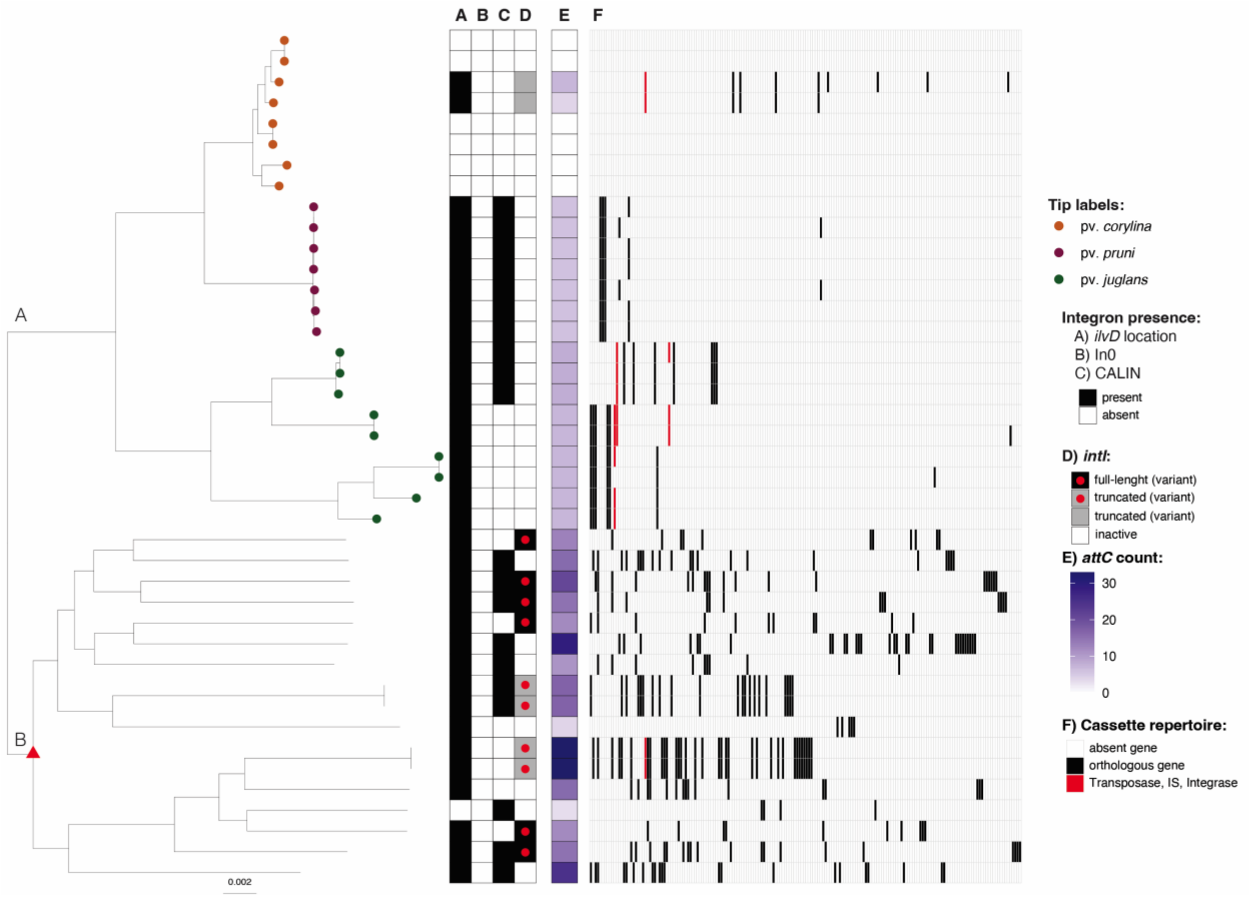


**Figure S3 – Integrons in *X. arboricola*.** The phylogeny of *X. arboricola* was built using Realphy, using GCA_000972745.1 as a reference genome. The tree was rooted with GCA_001908755.1 (*X. hortorum*) then the tip was removed from the tree. Tip labels show pathovars, which were assigned via literature search (Table S3). The red triangle indicates the probable acquisition of the *intI* variant. The scale bar indicates substitutions per site. A) Indicates the presence of a complete integron or *attC* sites in the *ilvD* locus, B) indicates the presence of an *intI* lacking cassettes (In0), C) indicates the presence of a CALINs (cluster of *attC* sites) not in the ilvD locus, D) indicates whether IntI is predicted to be functional (full length) (Table S3). E) indicates the number of *attC* sites (number of cassettes) in the genome. Panel F) represents the distribution of orthologous genes among the all cassettes carried by the corresponding isolate inferred with Proteinortho. Orthologous genes are given in decreasing order based on the number of strains within the species that carry them.

### *X. translucens*

*X*. *translucens* is the only species of group 1 that has more than 10 genomes in our dataset. *X. translucens* species can cause serious damage to cereals and to forage grasses, but also pistachio. Historically, two major groups have been distinguished within the species: the “translucens” and the “graminis” groups. Our dataset comprises genomes from clade Xt-I, which includes pvs. *hordei*, *translucens*, *undulosa*, and *secalis* (representing the “translucens” group), and clade Xt-III, which includes pvs. *arrhenatheri*, *graminis*, *phlei*, *phleipratensis*, and *poae* (the "graminis" group) (Goettelmann *et al.*, 2022). According to GTDB classification, clade Xt-II strains (*X. translucens* pv. *cerealis*) are a different species (*X. translucens_A)*.

Integron elements are present in 90.9% of the strains (Figure S4), but only 22.7% of the strains carry a full-length *intI*. Only the pv. *graminis* isolates (9.1% of the strains) lack both *attCs* and *intI*. As observed in other species, strains with the highest number of cassettes (15 or 16 *attCs*) contain an inactive *intI*. The cassette arrays in these strains harbor 74 orthologous genes, two of which encode transposases, with 33 genes (44.6%) identified as singletons. The most frequently observed cassettes, detected in 10 arrays across both Xt-I and Xt-III isolates, encode a hypothetical protein and a cassette encoding *symE*.

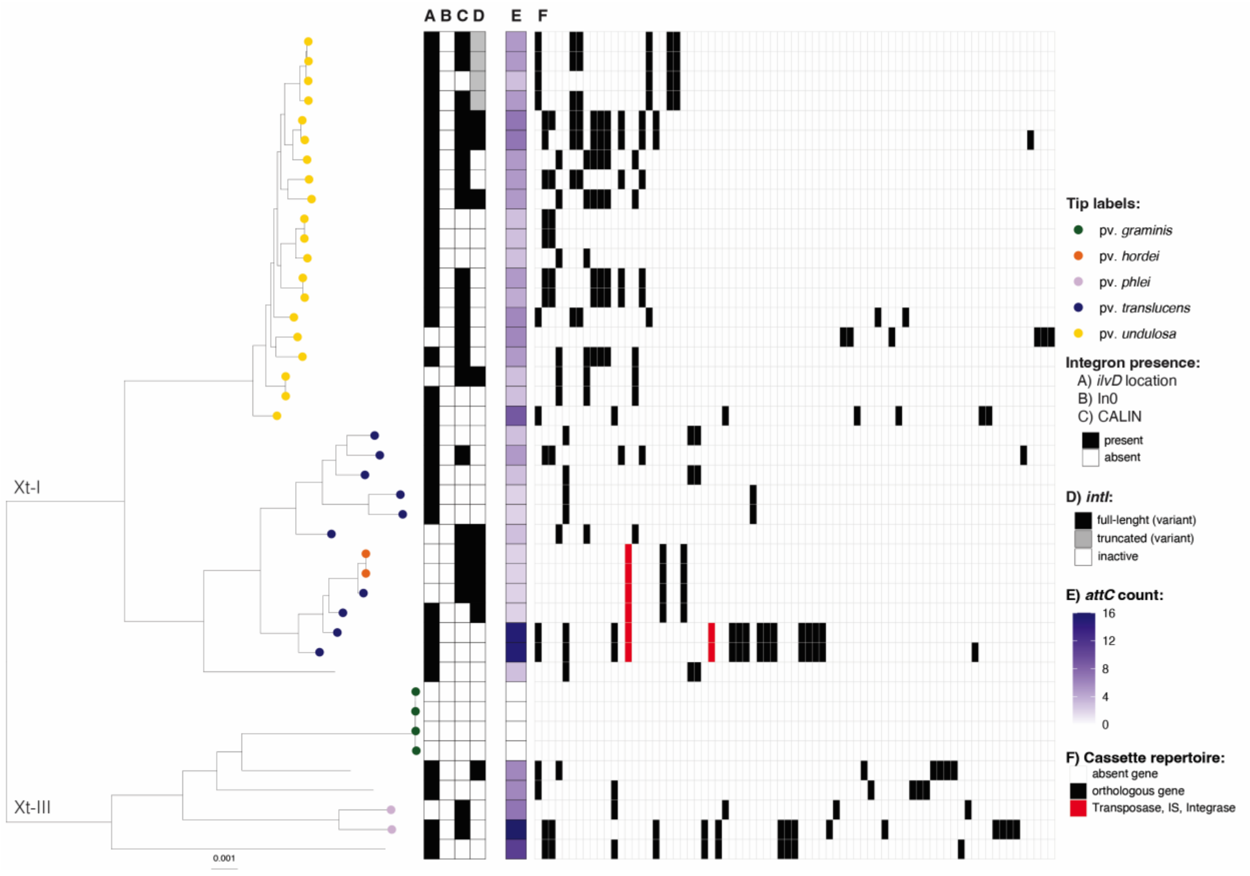


**Figure S4 – Integrons in *X. translucens*.** The phylogeny of *X. translucens* was built using Realphy, using GCA_001021935.1 as reference genome. The tree was rooted with GCA_000007145.1 (*X. campestris*) then the tip removed from the tree. Tip labels show pathovars, which were assigned via literature search (Table S3). The scale bar indicates substitutions per site. A) Indicates the presence of a complete integron or *attC* sites in the *ilvD* locus, B) inidicates the presence of an *intI* lacking cassettes (In0), C) indicate the presence of a CALINs (cluster of *attC* sites) not in the *ilvD* locus, D) indicates whether IntI is predicted to be functional (full length) (Table S3). E) indicates the number of *attC* sites (number of cassettes) in the genome. Panel F)

represents the distribution of orthologous genes among the all the cassettes carried by the corresponding isolate interfered with Proteinortho. Orthologous genes are ordered in decreasing order based on the number of strains within the species that carry them.

#### *X. euvesicatoria*

GTDB-Tk classifies as *X. euvesicatoria* isolates historically known as *X. euvesicatoria*, but also *X. axonopodis*, *X. perforans*, *X. campestris* pv. *fici* and pv. *olitorii*. The complete genomes of this species exhibit greater diversity compared to the species discussed above, largely because they do not represent an extensive expansion from a single pathovar; rather, the strains have been isolated from 21 different plant hosts.

Only one isolate (GCA_019444145.1) shows no evidence of integron structures, lacking both *attC* sites and *intI* (Figure S5). In contrast to the previously described species, *intI* appears to be more preserved, with 45.6% of the strains carrying a full-length *intI*. On average, the genomes contain 6.5 *attC* sites (ranging from 0 to 15), and the integron cassette arrays encode a total of 98 distinct orthologous genes, 10 of which are transposases or ISs. Approximately half of these genes (49 orthologous genes plus 2 transposases) are present as singletons.

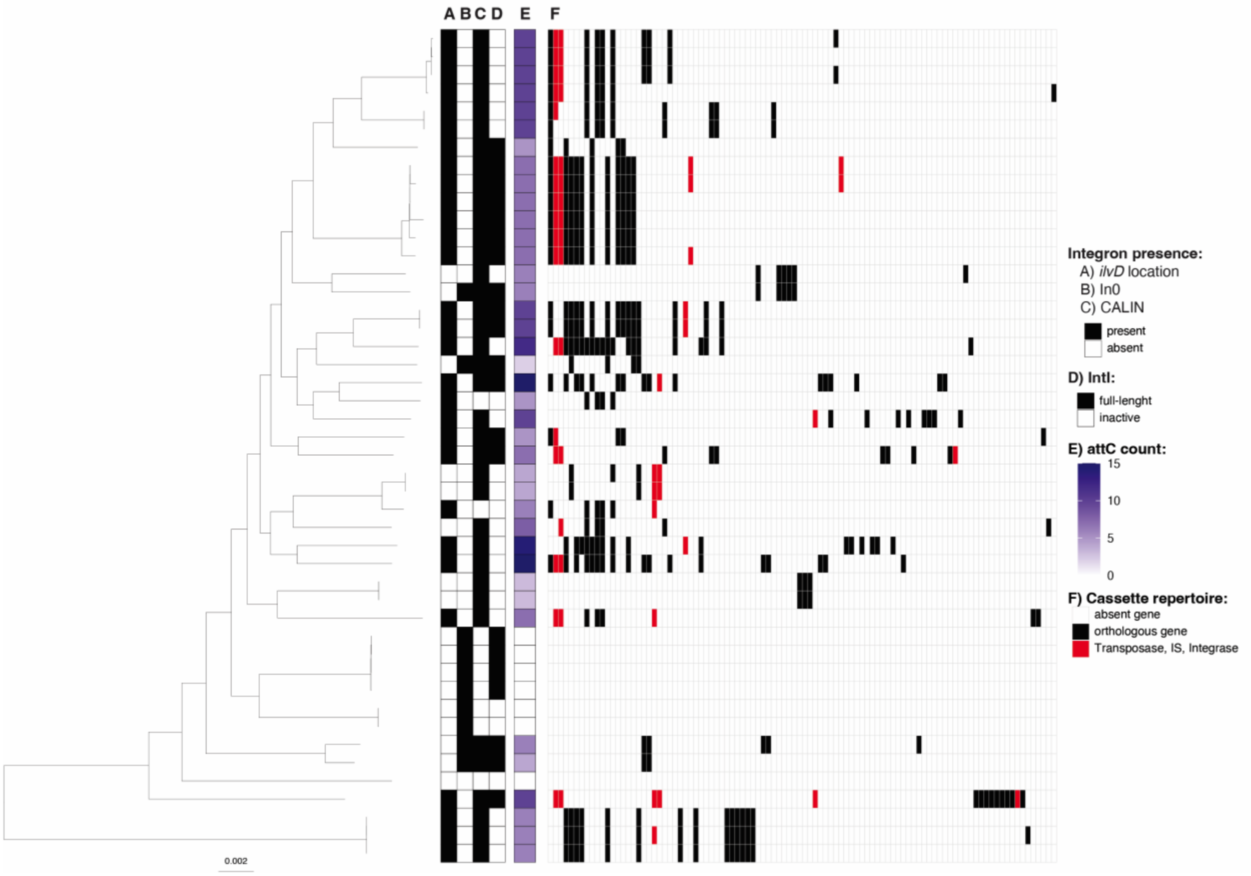

**Figure S5 – Integrons in *X. euvesicatoria*.** The phylogeny of *X. euvesicatoria* was built using Realphy, using GCA_000009165.1 as reference genome. The tree was rooted with GCA_002759095.2 (*X. phaseoli*) then the tip removed from the tree. The scale bar indicates substitutions per site. A) Indicates the presence of a complete integron or *attC* sites in the *ilvD* locus, B) inidicates the presence of an *intI* lacking cassettes (In0), C) indicate the presence of a CALINs (cluster of *attC* sites) not in the *ilvD* locus, D) indicates whether IntI is predicted to be functional (full length) (Table S3). E) indicates the number of *attC* sites (number of cassettes) in the genome. Panel F) represents the distribution of orthologous genes among the all the cassettes carried by the corresponding isolate interfered with Proteinortho. Orthologous genes are ordered in decreasing order based on the number of strains within the species that carry them.

### *X. oryzae*

*X. oryzae* is the causal agent of bacterial leaf blight (pv. *oryzae*) and bacterial leaf streak (pv. *oryzicola*) on rice. The strains used here include pv. *oryzicola*, pv. *oryzae* strains isolated in Asia and in Africa that form distinct phylogenetical groups, pv. *oryzae* strains recently introduced into Africa that are more similar to Asian strains (Schepler-Luu *et al.*, 2023), pv. *leersiae* strains, that are commonly pathogen of the pervasive weed species *Leersia hexandra* that frequently grows along rivers and canals surrounding rice paddies (Lang *et al.*, 2019), and X11-5A, a weakly pathogenic strain isolated in the USA in 1987 (Triplett *et al.*, 2011).

All *X. oryzae* isolates, except one, carry integrons, but only X11-5A carry a full-length *intI* (Figure S6). The pv. *oryzicola* isolates possess a truncated *intI* located at a locus that is not adjacent to *ilvD*. Although only three strains display a relatively high number of *attC* sites (ranging from 10 to 12 *attCs*), the average across genomes is less than two *attCs* per strain. Curiously, the strain with the full-length *intI* (X11-5A) harbors only 10 *attCs*, rather than the highest number observed. The integron cassette arrays in the species contain a relatively limited number of orthologous genes (61 in total), including 16 that encode transposases or ISs. Among these, 60.6%, comprising seven transposase or IS encoding genes, are singletons; however, of the 37 singletons identified, 18 are carried by a single pv. *leersiae* isolate (GCA_004319505.1). Although rice-associated *X. oryzae* strains generally exhibit greater genome plasticity compared to pv. *leersiae* (Lang *et al.*, 2019), the historical activity of their integrons appears to have been relatively higher.

Divergent cassette compositions were also observed among the pathovars. The Asian-like pv. *oryzae* strains possess cassettes encoding only transposases and ISs, whereas the African-like pv. *oryzae* strains harbor a cassette encoding a Transcription Activator-Like effector (TALe). TALes are effectors secreted via type III secretion systems into rice xylem. Once inside the host cell, they translocate to the nucleus where their unique domain of tandemly arranged 34-amino acid repeats mediates binding to specific promoter elements. Several TALes are known to induce host SWEET sucrose uniporter genes, thereby facilitating sucrose efflux from xylem parenchyma into the apoplasm at the infection sites. TALes may enhance diseases by targeting susceptibility genes, or may trigger a resistance response, and are therefore important for pathogen host range and virulence (Schepler-Luu *et al.*, 2023).

The TALe captured in the integron platform of the African-like pv. *oryzae* strains is also present in the Asian-like pv. *oryzae* strains with more than 91% amino acid identity and an identical number of long repeats. Curiously, the pv. *oryzicola* strains do not carry *attC* sites upstream of their truncated *intI*, yet they maintain an array of three TALes in the same locus. This observation suggests that either the TALe genes were originally acquired as integron cassettes and later fixed following *attC* sequence degradation, or that they were inserted at the locus through transposon-mediated mechanisms.

Collectively, these findings support the hypothesis that integron activity in *X. oryzae* was ancestrally active, followed by progressive inactivation concomitant with pathovar diversification.

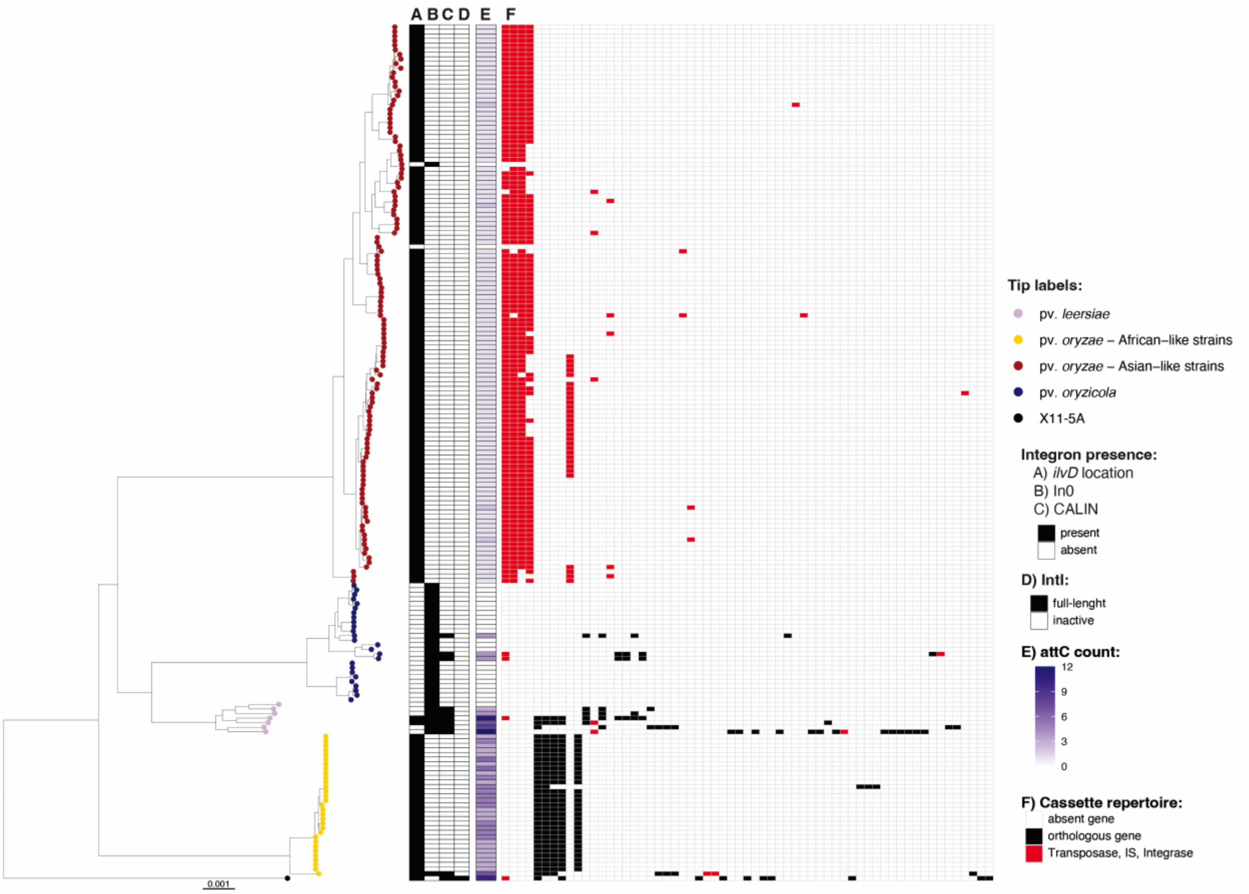


**Figure S6 – Integrons in *X. oryzae*.** The phylogeny of *X. oryzae* was built using Realphy, using GCA_000007385.1 as reference genome. The tree was rooted with GCA_002759095.2 (*X. phaseoli*) then the tip removed from the tree. Tip labels show pathovars, which were assigned via literature search (Table S3).

The scale bar indicates substitutions per site. A) Indicates the presence of a complete integron or *attC* sites in the *ilvD* locus, B) inidicates the presence of an *intI* lacking cassettes (In0), C) indicate the presence of a CALINs (cluster of *attC* sites) not in the *ilvD* locus, D) indicates whether IntI is predicted to be functional (full length). E) indicates the number of *attC* sites (number of cassettes) in the genome (Table S3). Panel F) represents the distribution of orthologous genes among the all the cassettes carried by the corresponding isolate interfered with Proteinortho. Orthologous genes are ordered in decreasing order based on the number of strains within the species that carry them.

#### *X. phaseoli*

The dataset analysed here predominantly comprises pathogenic strains of pv. *phaseoli* (Chen *et al.*, 2021), isolated from common bean, along with three strains from other hosts (sweet potato, cassava, and lace leaf). All isolates exhibited at least one integron signature (Figure S8). Notably, every *intI* gene detected in these strains is shorter than 700 bp, suggesting they are no longer functional. Within the pv. *phaseoli*, there is evidence that *intI* inactivation occurred twice, as the two basal strains (Figure S7) harbor an *intI* interrupted by a transposase distinct from those in the other pv. *phaseoli* isolates.

Despite the inactivation of *intI*, the isolate from lace leaf (GCA_008639345.1) and the isolate from sweet potato (GCA_041475645 .1) harbour 18 and 12 *attC*s respectively. However, the average number attC sites per genome in *X. phaseoli* is 2.5. A total of 26 orthologous genes were identified, five of which encode transposases or ISs. Among these, 38.5% (comprising nine orthologous genes and one transposase) were found to be singletons.

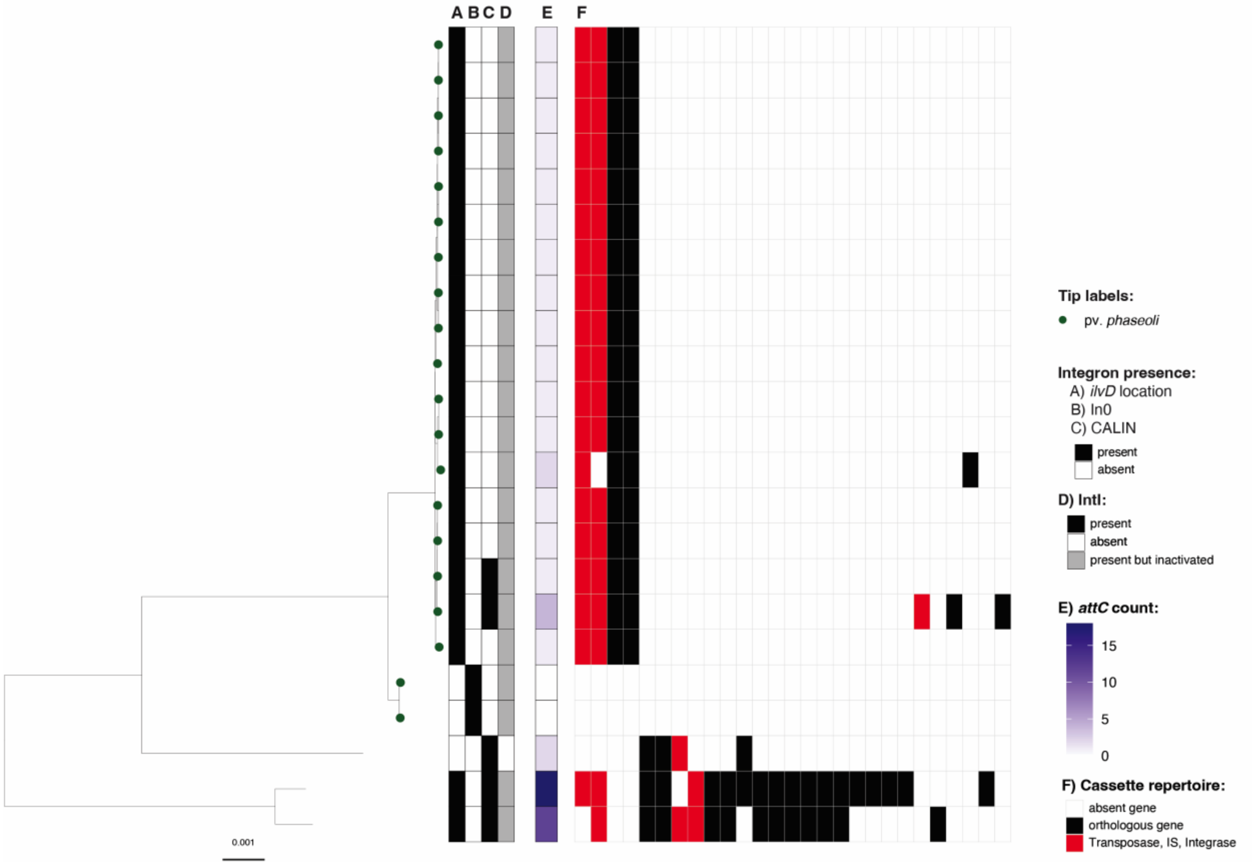


**Figure S7 – Integrons in *X. phaseoli*.** The phylogeny of *X. phaseoli* was built using Realphy, using GCA_008639345.1 as reference genome. The tree was rooted with GCA_000009165.1 (*X. euvesicatoria*) then the tip removed from the tree. Tip labels show pathovars, which were assigned via literature search (Table S3). The scale bar indicates substitutions per site. A) Indicates the presence of a complete integron or *attC* sites in the *ilvD* locus, B) inidicates the presence of *intI* lacking cassettes (In0), C) indicate the presence of a CALINs (cluster of *attC* sites) not in the *ilvD* locus, D) indicates whether IntI is predicted to be functional (full length) (Table S3). E) indicates the number of *attC* sites (number of cassettes) in the genome. Panel F) represents the distribution of orthologous genes among the all the cassettes carried by the corresponding isolate interfered with Proteinortho. Orthologous genes are ordered in decreasing order based on the number of strains within the species that carry them.

### *X. hortorum*

The pathovars of *X. hortorum* are responsible for causing bacterial spot and bacterial blight on a variety of host plants. The complete genomes analysed in this study derive from strains isolated from ivy (pv. *hederae*), dandelion (pv. *taraxaci*), lettuce (pv. *vitians*), and tomato (pv. *gardneri*) (Dia *et al.*, 2022).

All strains harbor CALINs and contain *attC* sites at the *ilvD* locus. However, only 42% of the isolates (5 out of 12) possess a full-length *intI* (figure S8). The genomes retain between 4 and 17 *attC* sites, with an average of 10.9 *attCs* per genome. Within the integron cassette arrays, 56 orthologous genes were identified, including 4 encoding transposases; despite the low number of genome analysed, 58.9% of the orthologues genes (33 genes, including 2 transposases) are present as singletons. The most commonly observed orthologous gene is *symE*, which is found in the integron arrays of all but two of the genomes.

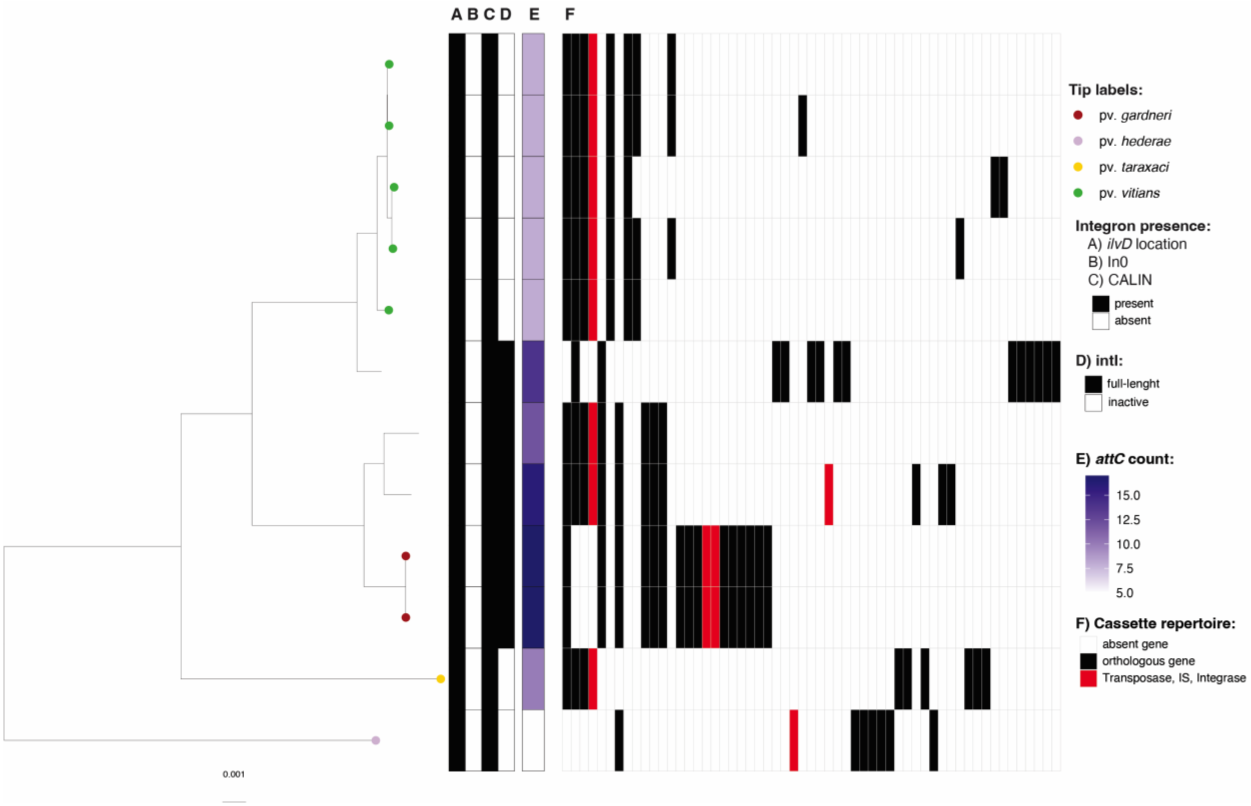


**Figure S8 – Integrons in *X. hortorum*.** The phylogeny of *X. hortorum* was built using Realphy, using GCA_002285515.1 as reference genome. The tree was rooted with GCA_905367715.1 (*X. arboricola*) then the tip removed from the tree. Tip labels show pathovars, which were assigned via literature search (Table S3). The scale bar indicates substitutions per site. A) Indicates the presence of a complete integron or *attC* sites in the *ilvD* locus, B) indicates the presence of an *intI* lacking cassettes (In0), C) indicate the presence of a CALINs (cluster of *attC* sites) not in the *ilvD* locus, D) indicates whether IntI is predicted to be functional (full length). E) indicates the number of *attC* sites (number of cassettes) in the genome (Table S3). Panel F) represents the distribution of orthologous genes among the all the cassettes carried by the corresponding isolate interfered with Proteinortho. Orthologous genes are ordered in decreasing order based on the number of strains within the species that carry them.

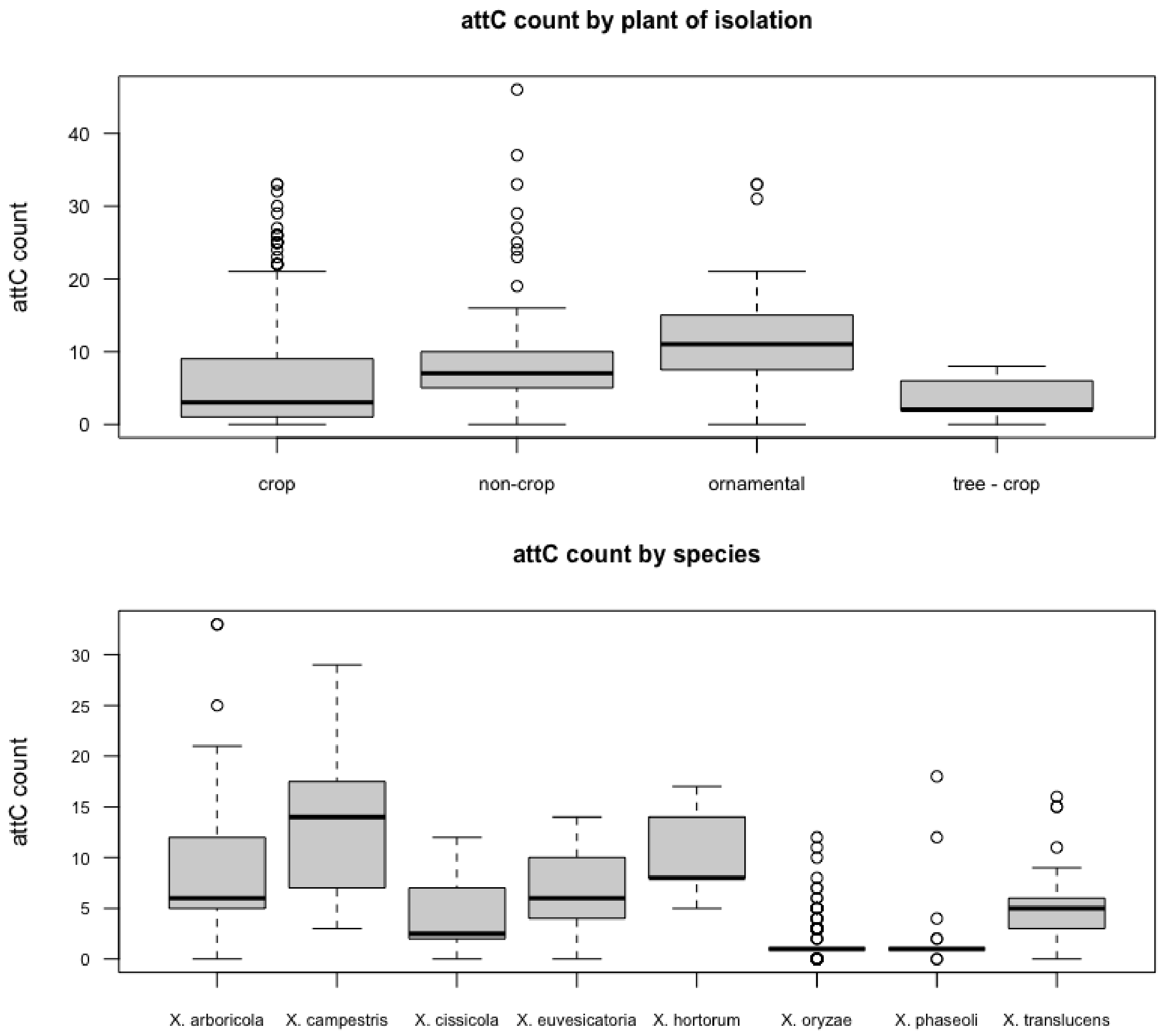


**Figure S9 – Integron gene cassette counts in *Xanthomonas*.** Boxplot of the number of *attC* sites per genome grouped by plant of isolation and by species (only species with more than 10 genomes are shown).

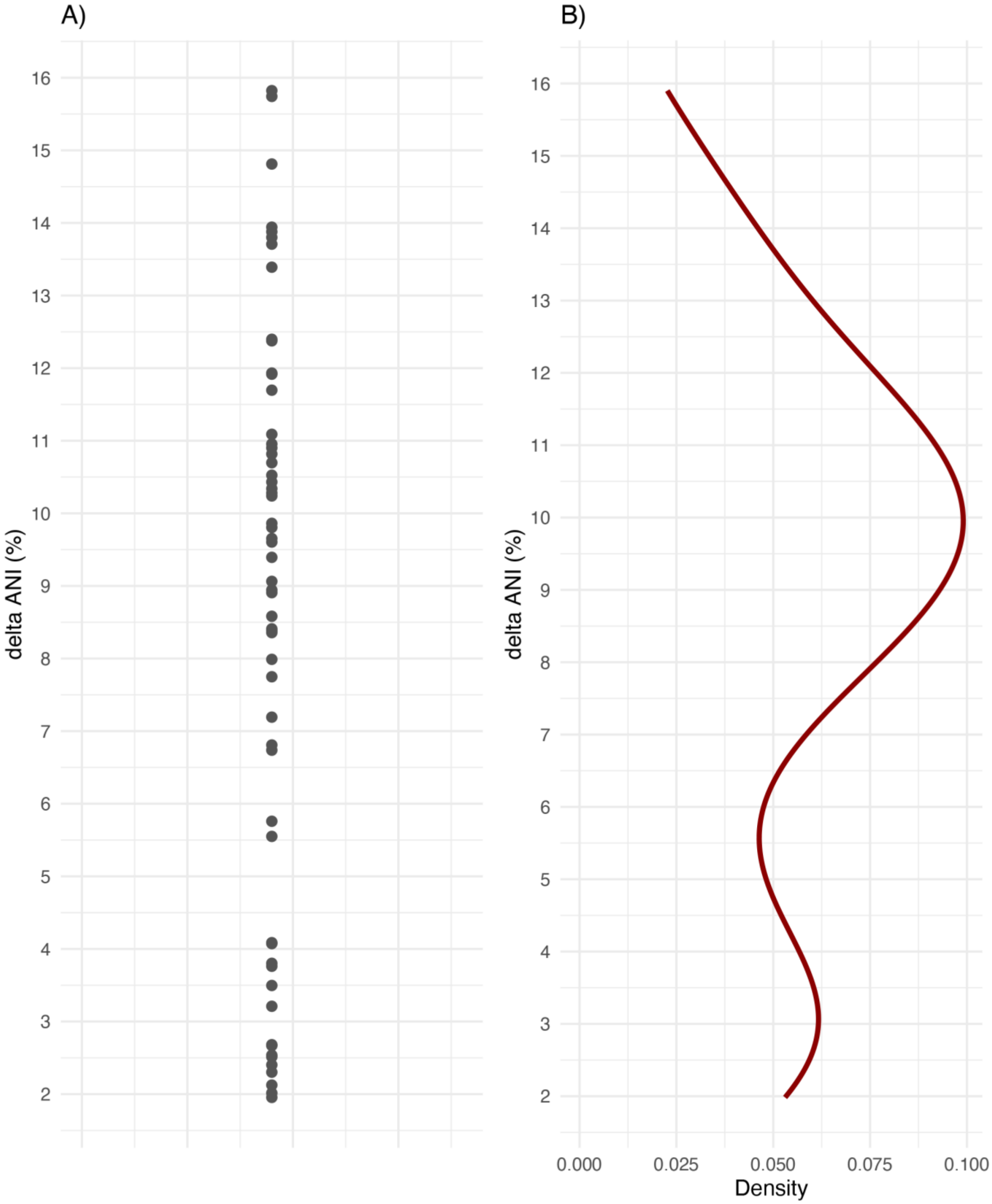


**Figure S10 – Delta ANI values for selected events of interspecies transfer of cassettes.** A) Dot plot and B) density plot of the cassette-to-genome ANI differences (delta ANI). In A), black dots represent individual data points, while in B), the curve shows the density estimate of the distribution.

### REFERENCES


Bansal K, Midha S, Kumar S & Patil PB (2017) Ecological and evolutionary insights into *Xanthomonas citri* pathovar diversity. *Applied and environmental microbiology* **83**: e02993-02916.

Chen NW, Ruh M, Darrasse A, Foucher J, Briand M, Costa J, Studholme DJ & Jacques MA (2021) Common bacterial blight of bean: a model of seed transmission and pathological convergence. *Molecular Plant Pathology* **22**: 1464-1480.

Delannoy E, Lyon B, Marmey P, Jalloul A, Daniel J, Montillet J, Essenberg M & Nicole M (2005) Resistance of cotton towards *Xanthomonas campestris* pv. *malvacearum*. *Annu Rev Phytopathol* **43**: 63-82.

Dia NC, Morinière L, Cottyn B, Bernal E, Jacobs JM, Koebnik R, Osdaghi E, Potnis N & Pothier JF (2022) *Xanthomonas* hortorum–beyond gardens: current taxonomy, genomics, and virulence repertoires. *Molecular plant pathology* **23**: 597-621.

Dubrow ZE, Carpenter SC, Carter ME, Grinage A, Gris C, Lauber E, Butchachas J, Jacobs JM, Smart CD & Tancos MA (2022) Cruciferous weed isolates of *Xanthomonas campestris* yield insight into pathovar genomic relationships and genetic determinants of host and tissue specificity. *Molecular Plant-Microbe Interactions* **35**: 791-802.

Fonseca NP, Patané JS, Varani AM, Felestrino ÉB, Caneschi WL, Sanchez AB, Cordeiro IF, Lemes CGdC, Assis RdAB & Garcia CCM (2019) Analyses of seven new genomes of *Xanthomonas citri* pv. *aurantifolii* strains, causative agents of citrus canker B and C, show a reduced repertoire of pathogenicity-related genes. *Frontiers in Microbiology* **10**: 2361.

Goettelmann F, Roman-Reyna V, Cunnac S, Jacobs JM, Bragard C, Studer B, Koebnik R & Kölliker R (2022) Complete genome assemblies of all *Xanthomonas translucens* pathotype strains reveal three genetically distinct clades. *Frontiers in Microbiology* **12**: 817815.

Lang JM, Pérez-Quintero AL, Koebnik R, DuCharme E, Sarra S, Doucoure H, Keita I, Ziegle J, Jacobs JM & Oliva R (2019) A pathovar of *Xanthomonas oryzae* infecting wild grasses provides insight into the evolution of pathogenicity in rice agroecosystems. *Frontiers in plant science* **10**: 507.

Patané JS, Martins J, Rangel LT, Belasque J, Digiampietri LA, Facincani AP, Ferreira RM, Jaciani FJ, Zhang Y & Varani AM (2019) Origin and diversification of *Xanthomonas citri* subsp. *citri* pathotypes revealed by inclusive phylogenomic, dating, and biogeographic analyses. *BMC genomics* **20**: 1-23.

Raga-Carbajal E, Espin G, Ayala M, Rodríguez-Salazar J & Pardo-López L (2022) Evaluation of a bacterial group 1 LEA protein as an enzyme protectant from stress-induced inactivation. *Applied Microbiology and Biotechnology* **106**: 5551-5562.

Rao MJ, Zuo H & Xu Q (2021) Genomic insights into citrus domestication and its important agronomic traits. *Plant Communications* **2**.

Schepler-Luu V, Sciallano C, Stiebner M, Ji C, Boulard G, Diallo A, Auguy F, Char SN, Arra Y & Schenstnyi K (2023) Genome editing of an African elite rice variety confers resistance against endemic and emerging *Xanthomonas oryzae* pv. *oryzae* strains. *Elife* **12**: e84864.

Triplett LR, Hamilton J, Buell C, Tisserat N, Verdier V, Zink F & Leach J (2011) Genomic analysis of *Xanthomonas oryzae* isolates from rice grown in the United States reveals substantial divergence from known *X. oryzae* pathovars. *Applied and Environmental Microbiology* **77**: 3930-3937.
